# Surfactant-Enhanced Hybridization Capture for Direct Sequence-Specific RNA Extraction from Plasma and Whole Blood

**DOI:** 10.64898/2026.10.06.757078

**Authors:** Wookyeom Kim, Shane Gilligan-Steinberg, Enos C. Kline, Ingrid Beck, Lisa M Frenkel, Barry Lutz

## Abstract

Direct recovery of low-abundance RNA from complex biological samples remains a major challenge for ultrasensitive detection. Sequence-specific hybridization capture enables selective enrichment, but recovery from crude samples is compromised by RNA degradation, protein interference, RNA secondary structure, and target sequence variation. Here, we address these limitations with a sodium dodecyl sulfate (SDS)-based formulation that not only lyses virions and protects RNA, but also directly accelerates hybridization kinetics, substantially reducing the need for thermal denaturation. This enabled direct RNA capture from crude plasma or whole blood with near-quantitative recovery, outperforming a widely used silica-based kit at low input. Without centrifugation or plasma separation, we detected HIV RNA by RT-qPCR at 25 copies/mL directly from whole blood. We further developed a free energy (ΔG)-guided algorithm for mutation-robust probe design, achieving >85% mean recovery across HIV-1 subtypes with a single probe. This work opens low-abundance RNA biomarkers in crude biofluids to ultrasensitive detection, addressing a major barrier to early diagnosis.

## Introduction

Nucleic acid detection is ubiquitous in healthcare and biology and provides a powerful approach for disease diagnosis, genetic studies, and fundamental biological research^1–9^. However, biological samples often contain high concentrations of interfering components that inhibit downstream molecular analysis^10,11^. Therefore, effective nucleic acid extraction that achieves both high recovery and high purity is critical for sensitive molecular analysis. Common nucleic acid extraction methods include phase partitioning (e.g., using phenol-chloroform), solid-phase reversible immobilization (SPRI) using PEG and salt, and solid-phase reversible nucleic acid adsorption (e.g., silica with high concentration chaotropic salts). Although these methods are convenient and well established, they non-selectively co-extract target and non-target nucleic acids from samples, which can mask detection of low-abundance targets^12,13^. Recovery from these methods also often declines at low target concentrations, and increasing sample input frequently fails to improve it, largely due to co-extracted PCR inhibitors that compromise downstream amplification^14^.

Low-abundance nucleic acid biomarkers are important candidates for clinical diagnostics^13,15–17^, but high levels of background nucleic acids in biological samples can overwhelm processes of purification and detection and undermine the biomarker detection^18^. In contrast, enrichment by sequence-specific hybridization to a DNA or RNA target can enable a substantial increase in the relative fraction of target nucleic acids recovered^19^. This capability is particularly important in scenarios where target nucleic acids are extremely scarce or obscured by highly abundant background nucleic acids. For example, our group achieved 84% clinical sensitivity using hybridization capture of tuberculosis cell-free DNA (cfDNA) from 10 mL urine, despite extremely short (∼40 nt) and low target DNA concentrations (<15 copies per mL median)^16,20,21^. Similarly, another study demonstrated that depleting platelet-derived RNA using a hybridization capture enabled the detection of tumor-derived cell-free RNA (cfRNA) with nearly a 50-fold improvement in sensitivity^15^. Similar benefits of hybridization-based enrichment have also been reported in wastewater surveillance^22^ and bloodstream infection diagnostics^23^. Together, these examples illustrate the broad utility of sequence-specific enrichment for recovering low-abundance nucleic acids from complex biological samples.

Nevertheless, sequence-specific RNA enrichment has not yet been widely adopted due to several limitations. First, achieving high recovery from long RNA targets remains challenging. While hybridization capture has been applied successfully to short RNAs such as miRNAs^24–26^, which are typically ∼20-24 nt, reported capture yields for whole viral RNA genomes have ranged from ∼7% to 40%^27,28^. The extensive secondary structure of long RNAs can reduce hybridization accessibility and slow hybridization kinetics, resulting in lower recovery or impractically long incubation times^29^. Heating and macromolecular crowding agents have each been shown to improve RNA hybridization efficiency, yet their quantitative contribution to RNA recovery in hybridization-based extraction remains unclear^30,31^. Second, direct RNA capture can be further complicated by the sample matrix itself. For example, abundant RNases in plasma samples can degrade RNA within minutes^32^, and non-specific adsorption of plasma proteins on magnetic beads can form a protein corona^33^, both of which contribute to reduced recovery^32,34^. Chaotropic salts, a common strategy for protecting RNA, can disrupt reagents used in hybridization capture (e.g., streptavidin-coated magnetic beads). Protein digestion does not provide immediate or complete protection from plasma RNase and requires multiple additional steps^35^. Third, hybridization efficiency is strongly impaired by sequence mismatches, which can lead to subtype-dependent differences in recovery and consequently reduce robustness across diverse variants of the target. Collectively, these limitations reduce efficiency, increase operational complexity, and complicate robust capture probe design.

Here, we quantitatively examined the effects of a pre-heating step and probe length on hybridization efficiency. We further show that a high concentration of an anionic surfactant not only protects RNA in complex biological matrices, including whole blood, but also directly accelerates hybridization kinetics. Based on these results, we established conditions that enabled near-quantitative recovery of long HIV RNA targets without detectable fragmentation. To address sequence variability, we developed a thermodynamic algorithm to identify capture probe binding sites predicted to tolerate sequence mismatches across subtypes. Using a single algorithm-designed probe, we demonstrated efficient extraction of HIV RNA, one of the most genetically diverse RNA viruses^36–38^, across multiple subtypes and clinical plasma samples. Together, these results establish a practical framework for sequence-specific extraction of long RNA from complex biofluids with broad applicability across targets and sample types.

## Materials and Methods

All experiments with heat-inactivated HIV and clinical plasma samples were conducted under BSL-2 conditions, while experiments involving cultured HIV were performed under BSL-2+ conditions, in accordance with approvals from the University of Washington Environmental Health & Safety (UW EH&S). Use of the de-identified remnant plasma specimens was determined to not involve human subjects by the University of Washington’s Human Subjects Institution Review Committee, and participant consent was therefore not required.

### Synthetic RNA template preparation (in vitro transcription)

To compare capture probe lengths and to assess RNA fragmentation after capture, a 266-nt RNA and a 4-kb RNA, respectively, were synthesized by in vitro transcription (IVT). For the 266-nt RNA, a DNA template containing a T7 promoter was obtained as a gBlock (IDT). For the 4-kb RNA, ∼9 kb genomic DNA extracted from the 8E5 cell line (ATCC, CRL-3617) served as the HIV DNA PCR template as described from the literature^39^. A ∼4-kb subfragment was then amplified from this product using Q5 High-Fidelity DNA polymerase (NEB, M0491), during which a T7 promoter sequence was introduced by PCR to generate the IVT template. The resulting fragment was purified from a 1% agarose gel using a QIAquick Gel Extraction Kit (QIAGEN, 28704). Primer sequences and PCR protocols are provided in the Supplementary Information.

For both templates, IVT was performed using Hi-T7 RNA Polymerase with the supplied buffer (NEB, M0658S) at 50 °C for 1 hr, followed by treatment with RNase-free DNase I (NEB, M0303S) for 15 min at 37 °C. The transcribed RNA was purified using the Monarch® Spin RNA Cleanup Kit (NEB, T2050S). The integrity of both RNAs was confirmed via an Agilent RNA ScreenTape assay on a 2200 TapeStation system (Agilent Technologies, 5067-5576, -5577, -5578). All reagents and buffers were used according to the manufacturer’s instructions. The 266-nt RNA was used solely for the capture probe length comparison experiment, and the 4-kb RNA was used on the same day of synthesis for the fragmentation test.

### Purified full-length HIV RNA template preparation

Heat-inactivated HIV stock of HIV type-1 strain IIIB was obtained through a customized order (ZeptoMetrix, 0801032CFHI). Viral RNA was isolated from the stock using the QIAamp Viral RNA Mini Kit (Qiagen, 52904) according to the manufacturer’s instructions. The purified RNA was aliquoted and preserved at −80 °C until further use. RNA concentration was determined with the GeneProof HIV-1 D PCR Kit (GeneProof, HIV1D/GP/0.25).

### EQAPOL Samples Extraction and Quantification

A panel of diverse HIV isolates was obtained from the EQAPOL repository^40^, representing major HIV-1 M group subtypes, circulating recombinant forms (CRFs), and unique recombinant forms (URFs), thereby reflecting the global genetic diversity of HIV. The isolates analyzed here were derived from cultured virus stocks originally established from HIV-positive plasma specimens. Viral RNA was extracted from 140 μL of culture supernatant using the QIAamp Viral RNA Mini Kit (QIAGEN, 52904) according to the manufacturer’s instructions and eluted in 60 μL of elution buffer. Extracted RNA was quantified using the GeneProof HIV Type 1 (HIV-1) D PCR Kit (GeneProof, HIV1D/GP/025). RNA samples were aliquoted and stored at −80 °C until use.

### HIV-positive patient’s plasma samples

A stock of HIV-positive plasma was made by pooling 20 de-identified remnant plasma specimens from HIV-infected individuals obtained from the University of Washington Clinical Virology Laboratory, Seattle, WA. The viral load of the pooled plasma stock was reported as 5.78 log_10_ c/mL, as determined by the University of Washington Clinical Virology Laboratory (quantified by Abbott RealTime HIV-1 VL assay, Abbott Molecular Diagnostics) and was further diluted with uninfected plasma obtained from an HIV-negative donor.

### Design of sequence-specific probes

Capture probes are DNA sequences that hybridize to a binding site in the target RNA. Each capture probe consists of a 5′ biotin, a 20-nt poly(A) spacer, a target-specific capture region, and a 3′ C3 spacer. The 5′ biotin provides a strong and stable complex with streptavidin on the magnetic bead surface, and the poly(A) spacer minimizes steric hindrance by distancing the hybridized target from the bead surface. The 3′ C3 spacer blocks the 3′ hydroxyl group, preventing polymerase extension when capture probes are released into the eluate for PCR analysis.

### Pre-immobilization of DNA capture probe on magnetic bead

Biotinylated single-stranded DNA (ssDNA) probes were synthesized by Integrated DNA Technologies (IDT). Probe sequences are listed in the Supplementary Table S1. Dynabeads MyOne Streptavidin C1 magnetic beads (Thermo Fisher Scientific, 65002) were washed twice with high-salt wash buffer (0.5 M NaCl, 25 mM EDTA, 10 mM Tris-HCl, 0.1% Tween-20) at a volume equal to the initial bead suspension. The beads were then incubated for 15 min with 40 pmol of biotinylated ssDNA probe per 20 µL of bead suspension in high-salt wash buffer. This was followed by washing steps: beads were washed with an equal volume of high-salt wash buffer to remove unbound probes and non-specifically hybridized DNA fragments, and then washed quickly with an equal volume of 30 mM NaOH to further remove non-specific binding. The beads were subsequently washed with an equal volume of 100 mM Tris-HCl (pH 8) to neutralize the pH. Finally, the beads were resuspended in storage buffer (low-EDTA TE buffer, 0.1 mM EDTA, with 0.1% Tween-20) at the same volume as the initial bead suspension and stored at 4 °C until use.

### Hybridization capture workflow

#### Sample preparation and hybridization

Unless otherwise noted, samples were prepared to a final volume of 1 mL. 250 µL HIV-negative human plasma, porcine whole blood, or PBS was mixed with hybridization buffer and SDS (final concentration: 0.5 M NaCl, 25 mM EDTA, 100 mM Tris-HCl pH 8, 0.1% Tween-20, 5% SDS). For SDS titration experiments, SDS was varied as indicated. To evaluate RNA recovery efficiency, dilution series of each RNA target were prepared in PBS. Purified RNA was spiked into plasma after SDS addition and mixing, whereas heat-inactivated HIV virion or HIV-1 positive plasma samples were spiked at the beginning of sample preparation to mimic the actual diagnostic workflow. Probe-immobilized magnetic beads (20 µL) were added, vortexed, incubated at 65 °C for 10 min unless otherwise indicated, and then rotated at room temperature for 30 min.

#### Bead separation, washing, and elution

Due to the high sample viscosity from 5% SDS, initial bead separation was incubated on the magnetic stand for >10 min, whereas subsequent captures in SDS-free wash buffers required only ∼1 min. Beads were washed twice with 700 µL high-salt wash buffer and once with 700 µL low-salt wash buffer (15 mM NaCl, 10 mM Tris-HCl). Captured RNA was eluted with 20 µL 30 mM NaOH and immediately neutralized by transfer into 2.2 µL neutralization buffer (85.1 mM Tris-HCl, 149 mM HCl), yielding a final eluate volume of 22.2 µL.

### RNA extraction from porcine whole blood

Porcine whole blood preserved with Alsever’s anticoagulant was purchased from LAMPIRE Biological Laboratories (catalog no. 7204903). To prepare spiked samples, HIV-positive plasma was serially diluted with PBS to achieve concentrations ranging from 10^1^ to 10^3^ copies per 5 µL. Each 5 µL of the dilution series was spiked into 125 µL porcine whole blood, which was then mixed with 125 µL of PBS and processed following the sequence-specific RNA extraction protocol described above. This resulted in a final hybridization volume of 1 mL, in which the original blood represented 12.5% (v/v).

### Scaled-up RNA extraction from porcine whole blood (1 mL, 25 copies)

A 5 µL aliquot of the HIV-positive plasma containing 25 copies of HIV RNA was prepared by dilution in PBS and spiked into 1 mL of porcine whole blood. The blood sample was incubated at room temperature for 5 min. The 1 mL whole blood was then transferred to an Eppendorf LoBind 15 mL tube and mixed with 2.08 mL of hybridization buffer and 2.92 mL of PBS. 2 mL of SDS was added to achieve a final concentration of 5%, resulting in a final volume of 8 mL. A total of 50 µL of capture probe-immobilized magnetic beads were added to the mixture. The tube was placed on a hot plate and heated to 70 °C for 5 min. To improve heat transfer, 2 mL of water was added to the aluminum heating block wells. During heating, the tube was manually inverted every 2 min and reheated on the heating block for a total heating time of 15 min. Following the heating step, the tube was continuously rotated at room temperature for 15 min using a rotisserie mixer. Magnetic beads were collected using a custom magnetic stand assembled with neodymium block magnets (K&J Magnetics, Inc., BX088-N52) designed for 15 mL tubes (Supplementary Figure S1). After placement on the magnetic stand for 10 min to allow bead collection, the supernatant was removed using a serological pipette. The beads were resuspended in 1 mL of high-salt washing buffer, and the suspension was transferred to a 1.5 mL Eppendorf LoBind tube. All subsequent extraction steps were performed identically to the extraction protocol described above.

### Recovery rate calculation using RT-qPCR

One-step RT-qPCR was performed using OneTaq Hot Start DNA Polymerase (NEB) and WarmStart RTx Reverse Transcriptase (NEB) on a CFX96 Real-Time PCR System. Specific reaction compositions and cycling conditions, including primer sequences, are detailed in the Supplementary Information. Each PCR amplification region was located within ∼200 nt of the capture region to minimize fragmentation-related bias. Standard curves spanned 10¹ to 10⁴ copies per reaction for 10³-copy inputs and 10² to 10⁵ for 10⁴-copy inputs, with the range set by the available stock concentration so that all measured values fell within the curve. For the subtype panel, 10⁴ copies were spiked, and a separate standard curve from 10³ to 10⁵ copies was generated for each subtype. To ensure identical reaction conditions between the standard curve and extracted RNA samples, both reactions contained the same proportions of master mix, elution buffer, and total volume. Specifically, 22.8 µL of master mix was combined with 22.2 µL of eluate and 5 µL of TLE buffer for extracted samples, or with 22.2 µL of clean neutralized elution buffer and 5 µL of TLE buffer containing the RNA dilution series for quantification standards. Even though we confirmed that the neutralized buffer itself did not affect RT-qPCR Cq values, identical reaction compositions were maintained between the standard curve and extracted samples. The dilution series was stored at 4 °C during the capture process. Standard curve and experimental samples were subsequently amplified in parallel, and RNA quantities were determined based on the resulting standard curve. The recovery rate was calculated as (measured RNA copy number ÷ input copy number) × 100.

### Thermodynamic Profiling of the Synthetic RNA Target

To evaluate the fundamental thermodynamic interplay between target structure and probe hybridization, in silico profiling was first performed on a 266-nt synthetic RNA target. The structural accessibility of the target sequence was predicted using the RNAplfold algorithm from the ViennaRNA package (version 2.6.4)^41^. For each binding site, we calculated the probability that a contiguous 10-nt window was completely unpaired, and then the mean of these probabilities within the capture probe binding site was used to represent its accessibility (P_unpaired_). Concurrently, the bimolecular hybridization fraction between the target RNA and complementary DNA capture probes of varying lengths was calculated using a nearest-neighbor thermodynamic model for RNA–DNA hybrids, based on the parameters reported by Sugimoto et al. (1995)^42^, under 1 M Na⁺ conditions. Both folding and hybridization simulations were computed across a continuous temperature gradient (e.g., 25 °C to 100 °C at 5 °C increments) under a 1.0 M Na⁺ concentration with target and probe strand concentrations of 1 × 10⁻^17^ M and 4 × 10⁻^8^ M, respectively. For each temperature, standard-state duplex free energies were converted to equilibrium constants and the equilibrium duplex concentration was obtained from a 1:1 mass-balance model, with the hybridization fraction reported as [RNA-probe complex]/[Initial target RNA].

### Global Accessibility Landscape of the Full-Length HIV-1 Genome

A spatial thermodynamic analysis was performed on the full-length HIV-1 consensus sequence. Degenerate International Union of Pure and Applied Chemistry (IUPAC) codes and unresolved bases (’N’, ’?’) were randomly substituted with canonical bases (A, C, G, U). Targets containing ≥50% undefined bases were excluded from the analysis. The genome was scanned using a sliding window at 10-nt intervals. At each coordinate, the local target sequence was extracted with 150-nt flanking extensions at both ends to preserve structural context. Local accessibility was calculated as the mean P_unpaired_ across the capture region. For this genome-wide analysis, calculations were performed at 0.5 M Na⁺ to match the experimental conditions, at 25 °C and 65 °C.

### Plasma gelation test

Commercially available human plasma, used only for a gelation test, was obtained from Fisher Scientific (Cat. No.: 50-203-6372). Plasma samples were mixed with hybridization buffer containing increasing concentrations of SDS (Fisher Scientific, BP1311-200) from 0% to 9.8%. The mixtures were incubated at either 65 °C or 95 °C for 10 min. Sample turbidity was measured at 600 nm using a Biochrom WPA CO8000 spectrophotometer. The mixture maintained at room temperature was used as the control for turbidity measurement.

### SDS mechanism experiments

The *env*-probe sequence and RT-qPCR protocol are described in Supplementary Table S1In no-SDS conditions, SDS was replaced with an equal volume of water. All other steps followed the hybridization capture workflow described above. To determine at which step SDS acts, a dummy high-salt wash step was added before the standard washing steps in all groups so that the number of wash steps was identical across conditions, and three conditions were compared: 5% SDS during hybridization only, 5% SDS in the dummy wash only, and no SDS throughout. Separately, to test whether the effect was specific to SDS, 20% Triton X-100 and 15% Ficoll 400 (Fisher Scientific, AAB2209509) stock solutions were prepared, and each additive was substituted for SDS at a final concentration of 5% during hybridization.

### Capture probe design using algorithms

HIV sequence alignments were obtained from the Los Alamos National Laboratory (LANL) HIV Database (HIV-1/SIVcpz, Region: GENOME, Subtype: M group with CRFs, Year: 2021). The probe design algorithm was initially implemented in MATLAB and later translated into Python to incorporate the NUPACK module^43^. The algorithm accepts user-defined parameters. In this work, we used the following parameters. (Probe length: 70-nt, mismatch tolerance: 9, ΔG cutoff: -40 kcal/mol, GC content range: 40∼65%, monovalent ion concentration: 0.5 M, temperature: 65 °C, probe and target concentration: 4*10^-8^ and 1.7*10^-17^, and candidate region range: 0.9)

A consensus sequence was generated by selecting the majority nucleotide at each alignment position. The algorithm then scanned the consensus with a sliding window equal to the probe length, shifting by one nucleotide at a time. At each window position, the Hamming distance between the consensus subsequence and each aligned target subsequence was calculated. Insertions or deletions in the alignment were treated as mismatches, consistent with Hamming-based mismatch scoring. A target sequence was considered to satisfy the mismatch criterion if its Hamming distance within the window was less than or equal to the user-defined mismatch tolerance. Candidate probe regions were first filtered by GC content, excluding windows outside the range of 40% to 65% to avoid extremely weak (low GC) or overly stable and self-structured (high GC) probes. The remaining windows were ranked in descending order based on the number of target sequences that satisfied the mismatch cutoff. From among the highest-ranked regions, those with values of at least 90% of the maximum were selected for thermodynamic evaluation. The 90% threshold is a user-defined reduction step chosen empirically to retain all major peaks in the mismatch-based distribution while excluding regions that perform poorly and do not require full ΔG evaluation. For each retained candidate, hybridization Gibbs free energies (ΔG) were calculated between the candidate probe and all target sequences using the NUPACK Python module (Complex Analysis). To improve computational efficiency, the algorithm groups identical sequences among the targets and performs the NUPACK calculation only once for each unique sequence.

When IUPAC ambiguity codes were present in either the probe or target, the ambiguity code was replaced with one of the corresponding canonical nucleotides when evaluating ΔG. A target was considered capturable if the predicted ΔG was below the user-defined cutoff. Candidate regions were again ranked by the number of capturable target sequences. In parallel, accessibility was calculated across all target regions in the consensus sequence using the method described above. The final output table contained probe position, probe sequence, mean, minimum, and maximum ΔG values, ΔG standard deviation, the number of targets satisfying mismatch and ΔG criteria, overall predicted capture coverage, unpaired probability, and accessibility percentile. Accessibility percentile indicates how high the accessibility of a given region is relative to that of the entire genome. Users may select single or multiple probes depending on coverage and redundancy requirements. The computational algorithm for probe design is available at GitHub repository (https://github.com/WookyeomKim/RNA-Capture-Probe-Designer)

### Establishing ΔG threshold for RNA capture

A 5′-biotinylated ssDNA capture probe targeting the *gag* region was synthesized by IDT and intentionally designed to include mismatches among the EQAPOL panel sequences. Six HIV variants from the EQAPOL panel were selected to span a wide range of mismatch counts and distributions within the probe-binding site. For each target RNA, hybridization capture in PBS (n = 3) was performed and analyzed in parallel with a matched standard curve generated from purified HIV RNA of the same sequence (n = 3). Using a sequence-matched standard curve for each variant minimizes RT–qPCR bias arising from sequence-dependent amplification efficiencies, allowing recovery to be compared across variants. The hybridization free energy (ΔG) between the probe and target RNA was computed using the NUPACK complex analysis module under the same ionic and temperature conditions described above. Because NUPACK lacks RNA–DNA hybrid thermodynamic parameters, ΔG values were calculated using the DNA/DNA model. However, because the algorithm also uses the same DNA/DNA model, the ΔG cutoff is still compatible with the computation and remains predictive for capture performance. Probe and target sequences are provided in the Supplementary Information.

### Simulation of ΔG variation in 70-nt DNA duplexes

Simulation work was conducted in MATLAB by interfacing with the NUPACK Python module. Random pairs of 70-nt DNA sequences were generated with GC contents constrained between 40% and 65%. Increasing numbers of mismatches were introduced into one of the strands. The hybridization free energy (ΔG) between the two strands was calculated using the NUPACK complex analysis function (under the same conditions described above). This process was repeated 5,000 times for each mismatch number, starting from the generation of random DNA pairs. The variation in ΔG was plotted in Figure 6C.

### Comparison to the QIAamp Viral RNA Mini kit

HIV-positive plasma was diluted in PBS and spiked into HIV-negative donor plasma to give 0, 30, 300, and 3,000 copies of HIV RNA per 140 μL. For RT-qPCR, 20 μL of 60 μL eluate was used, corresponding to 10, 100, and 1,000 copies per reaction. For hybridization RNA capture, the procedure described above was followed, adjusting the final volume to 560 μL (equivalent to 140 μL of plasma) to match the input volume used in the QIAamp kit. RNA extraction with the QIAamp kit was performed according to the manufacturer’s protocol including all recommended steps. Although hybridization capture can concentrate RNA by reducing the elution volume to 22.5 μL, the eluate was diluted to 60 μL with TLE buffer to match the QIAamp kit’s recommended final eluate volume for fair comparison. A 20 μL aliquot of each eluate was used as input for a 50 μL RT-qPCR reaction. Both extraction methods were performed in parallel using the same dilutions of plasma samples, and all extracted RNA, along with purified HIV RNA, was analyzed by RT-qPCR on the same CFX instrument to calculate recovery rates.

### Fragmentation analysis

To assess whether the capture process fragments long RNA targets, 4 μL of truncated 4-kb synthetic HIV RNA, containing approximately 21 ng of RNA, were spiked into 200 μL of PBS. For control samples, 4 μL of the synthetic RNA stock was mixed separately with 18.2 μL of TLE buffer or neutralized elution buffer prepared by mixing 20mM NaOH and neutralization buffer at a 20:2.2 volumetric ratio to yield a near neutral pH (approximately 8∼9). 2 μL of each control was loaded directly onto the gel to assess whether ionic strength differences affected RNA migration. Two bead controls were included. The negative control contained capture probe–immobilized magnetic beads without target RNA, whereas the bare-bead control contained streptavidin magnetic beads without capture probes and target RNA. Hybridization capture was performed as described above. 2 μL of each eluate was analyzed by automated gel electrophoresis using the TapeStation 2200 system (Agilent Technologies) with High Sensitivity RNA ScreenTape (Agilent, 5067-5579, 5067-5580, 5067-5581). RNA from both the control and captured samples was quantified using the TapeStation software.

### Statistical analysis

All experiments were performed in triplicate unless otherwise stated. Data are expressed as mean ± SD. Two groups were compared by Student’s t-test, multiple groups by one-way or two-way ANOVA with Tukey’s post-hoc test, and ordered concentrations by the Jonckheere-Terpstra trend test, as indicated in the corresponding figure legends. Statistical significance was defined as follows: * p < 0.05, ** p < 0.01, *** p < 0.001, **** p < 0.0001.

## RESULTS AND DISCUSSION

### Sequence-specific RNA extraction via hybridization capture

Figure 1 illustrates the overall hybridization capture workflow. Streptavidin-coated magnetic beads pre-immobilized with biotinylated capture probes are directly added to plasma or blood samples in the presence of hybridization buffer. Following incubation with pre-heating and subsequent cooling to room temperature, the capture probes hybridize with target RNA. Washing steps remove plasma or blood components and nonspecific nucleic acids that could inhibit downstream amplification. Captured RNA is then quickly eluted at high pH, which disrupts hydrogen bonds between the probe and target, and the eluate is immediately neutralized for one-step RT-qPCR analysis. We used this system to investigate the effects of capture probe length, pre-heating, and buffer composition on hybridization efficiency, the stability of capture in crude plasma and whole blood, and the design of probes that tolerate target sequence variation.

**Figure 1.**
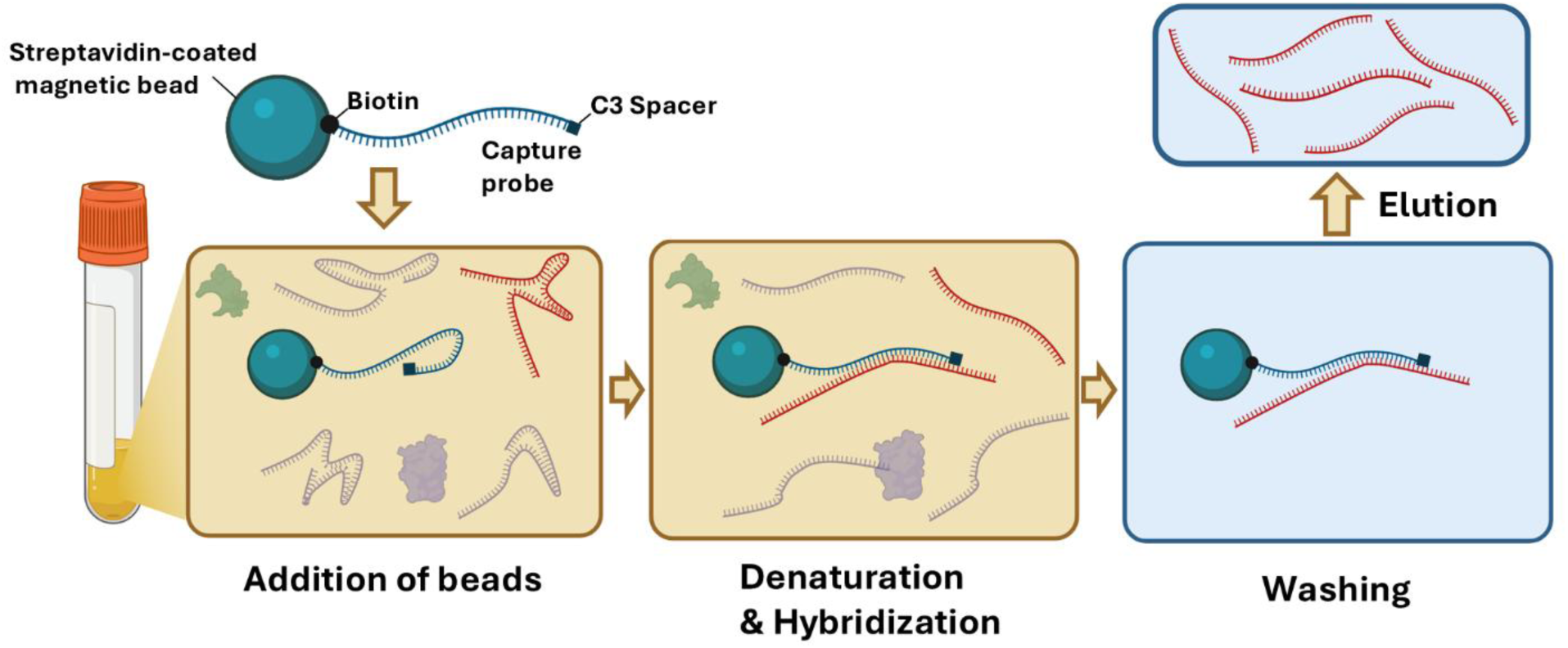
Illustration of sequence-specific RNA extraction via hybridization capture. Capture probes immobilized on magnetic beads hybridize with target RNA in plasma or whole blood, enabling selective binding, washing, and concentration of the target. (Created with BioRender.com)

### Effect of Heat on RNA Hybridization Efficiency

Nucleic acid secondary structure strongly impedes hybridization kinetics, leading to poor hybridization capture efficiency^44^. This effect is particularly pronounced for RNA targets due to their strong intramolecular folding^45,46^. Heating can relieve this barrier by unfolding the target^31,47^, but its contribution to recovery in bead-based capture of long RNA, and its dependence on probe length, remains unclear. To evaluate this effect, we tested capture efficiency from PBS using a 266-nt RNA fragment and capture probes of varying lengths from 19-nt to 120-nt, both with and without heating (Fig. 2A-C). Capture without pre-heating resulted in only 20–30% RNA recovery regardless of probe length (Fig. 2B). When samples and capture probes were pre-heated to 65 °C followed by passive cooling to room temperature, recovery increased for all probe lengths (Fig. 2C). Recovery with probes of 40-nt and longer was high (∼80-100%), while recovery with the 19-nt probe was improved by heating but remained low. This finding suggests the presence of substantial secondary structure within the target RNA, which is qualitatively illustrated by the NUPACK-predicted minimum free energy structure shown in Supplementary Figs. S2-S3.

**Figure 2.**
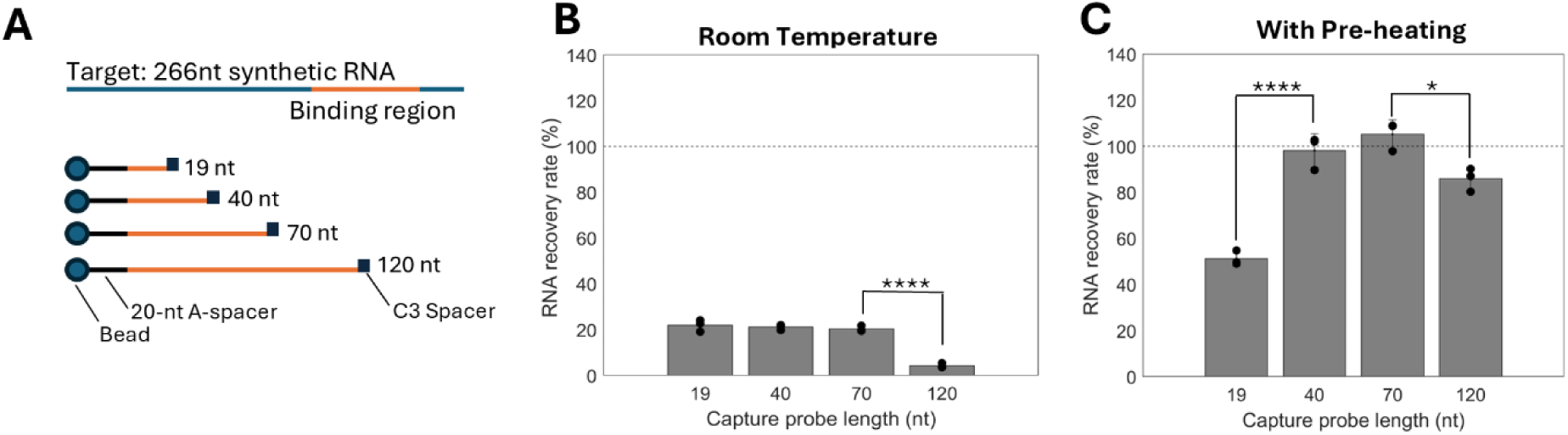
Influence of capture probe length and pre-heating on RNA extraction efficiency from PBS without SDS. (A) Illustration of the extraction experiment using a 266-nt synthetic RNA target (10^4^ copies) and capture probes of varying lengths. (B–C) RNA recovery rate across capture probe lengths at room temperature (B) or after pre-heating at 65 °C for 10 mins, followed by cooling to room temperature (C). All quantitative data are presented as mean ± SD (n = 3). Statistical comparisons were performed using one-way ANOVA with Tukey’s HSD post-hoc test.

The reduced capture efficiency of the 19-nt probe likely stems from its low melting temperature, which delays hybridization until cooling and forces the probe to compete with the target’s secondary structure (Supplementary Fig. S4). Although pre-heating substantially increases target RNA accessibility, subsequent cooling induces rapid intramolecular target refolding that outpaces short-probe binding. In contrast, longer probes (≥40 nt) remain duplexed at high temperatures (65 °C) and efficiently capture the target without secondary structure competition. Recovery declined again at 120-nt, possibly due to probe self-structure. We selected 70-nt because longer probes retain more thermodynamic margin per mismatch, which is advantageous for genetically diverse targets (see Algorithm-guided capture probe design).

### Effect of SDS on Viral Lysis, Plasma Stability, and RNA Preservation

Efficient RNA capture from plasma requires release from viral capsids or extracellular vesicles while protecting against RNA degradation by abundant RNases. In addition, elevated temperature is required to achieve efficient hybridization, but heating induces thermal gelation of plasma proteins. To address these challenges, we employed SDS, an anionic surfactant that denatures proteins, disrupts protein– protein interactions, and rapidly inactivates RNases^48^.

#### Viral lysis and temperature limit

HIV RNA is encapsulated within a viral capsid and lipid envelope. To evaluate the lytic performance of SDS, heat-inactivated HIV virions were spiked into PBS, and the RNA was recovered using the hybridization capture method (Fig. 3A). In the absence of both heating and SDS, RNA recovery was only ∼40%. While heating alone at 65 °C increased recovery to ∼80%, the addition of 1% SDS further enhanced recovery to near 100%. We observed a substantial decrease in RNA recovery when heating at 80 °C in 1% SDS, and reduced capture efficiency was also observed at 90 °C in the absence of SDS in our previous work, where it was rescued by dual biotin labeling^16^. Temperature alone can therefore destabilize the streptavidin–biotin complex, consistent with reported thermal and SDS-induced transitions of streptavidin^49,50^. Nevertheless, capture remained efficient at 65 °C, identifying this as a practical balance between viral lysis and capture stability.

**Figure 3.**
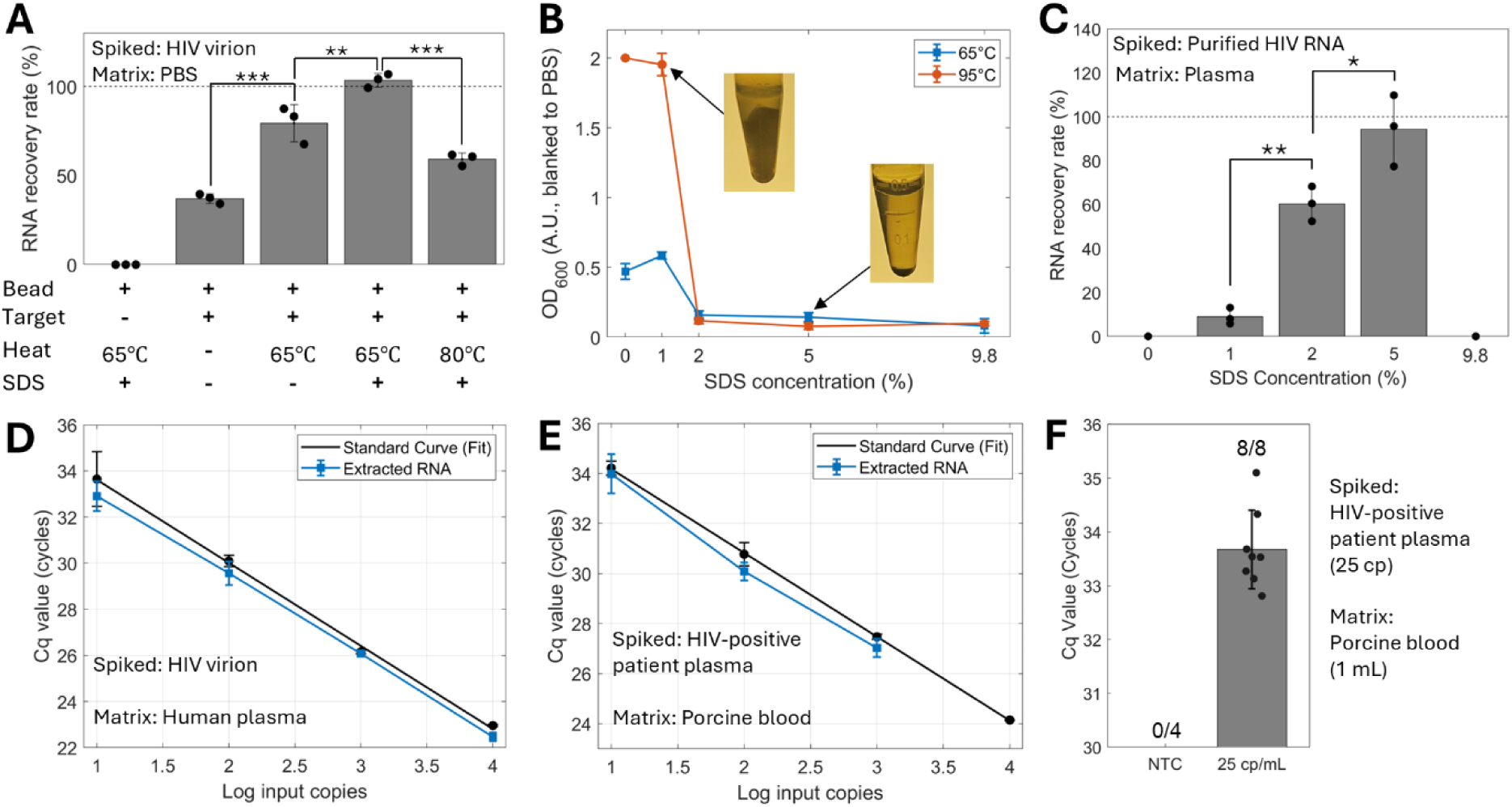
Effect of heating and SDS on plasma and blood stabilization and RNA recovery using a 70-nt capture probe targeting HIV RNA. (A) RNA recovery from heat-inactivated HIV virion (10^4^ copies) in PBS under different heating conditions, with or without 1% SDS. Statistical comparisons among the four target-containing conditions were performed by one-way ANOVA with Tukey’s HSD post-hoc test. (B) Turbidity of plasma mixture heated at 65 °C or 95 °C with increasing SDS concentrations, measured by absorbance at 600 nm. (C) Recovery of purified HIV RNA (10^4^ copies) spiked into plasma at increasing SDS concentrations. All conditions contained 25 mM EDTA and 100 mM Tris-HCl, pH 8. No amplification was detected at 0% or 9.8% SDS. Statistical comparisons (one-way ANOVA with Tukey’s HSD post-hoc test) were performed among the remaining groups (1, 2, 5%). (D, E) Recovery from heat-inactivated HIV virions in plasma and HIV-positive patient plasma in porcine whole blood at varying input copies using 5% SDS, quantified by standard curves (PCR efficiency, 90% and 98%, respectively). (F) Detection of 25 copies of HIV RNA from patient plasma spiked into 1 mL porcine whole blood, shown by Cq comparison between no-template controls (NTC, n = 4) and 25-copy samples (n = 8). Quantitative data are presented as mean ± SD (n = 3), unless otherwise indicated.

#### Plasma Stabilization

Thermal gelation of plasma emerged as a major impediment to direct genome extraction in our protocol. Plasma proteins lose structural stability above physiological temperature and substantially denature around 60–65 °C^51,52^, leading to protein–protein interactions and aggregation. Nonionic surfactants such as Tween-20 and Triton X-100 failed to prevent gelation even at concentrations above 9.8% (data not shown). We therefore tested SDS, adding increasing concentrations to the plasma mixture followed by heating at 65 °C or 95 °C for 10 min (Fig. 3B). At 65 °C, samples containing 0% or 1% SDS formed gels, resulting in an approximately 4-fold increase in turbidity. Unexpectedly, 1% SDS produced higher turbidity than the 0% condition (∼24% increase). This effect may reflect partial protein unfolding. At insufficient surfactant concentrations, SDS can expose hydrophobic regions without fully shielding them, potentially promoting intermolecular interactions^53^. In contrast, SDS concentrations above 2% prevented heat-induced plasma gelation even at 95 °C. However, 9.8% SDS, although suppressing gelation, rendered the mixture too viscous and was therefore not ideal for bead extraction.

#### RNA preservation

RNase inactivation is essential for high RNA recovery, as plasma RNases rapidly degrade RNA. Although chaotropic salts such as GuSCN are commonly used^54,55^, high concentrations (≥4 M) compromised bead surface properties and induced aggregation (Supplementary Fig. S5). Alternatively, SDS can fully inactivate RNase at concentrations as low as 0.5% by unfolding RNase structure^35^. Although this inactivation is reversible, RNases are removed during subsequent washing steps in our protocol.

We evaluated whether SDS protects RNA in plasma while remaining compatible with bead-based hybridization capture. Purified HIV RNA was spiked after SDS was mixed with the plasma and hybridization buffer. Despite the presence of 100 mM Tris-HCl and 25 mM EDTA in all conditions, no signal was detected without SDS, indicating that these components alone were insufficient. In contrast, 1% SDS yielded 8% recovery, increasing to 94% at 5% SDS (Fig. 3C). At 9.8% SDS, the bead extraction failed due to increased viscosity and bead aggregation. We also confirmed that SDS carryover did not affect RT-qPCR, as Cq values were unchanged between PBS and 5% SDS samples (Supplementary Fig. S6).

### Evaluation of Analytical Sensitivity

Achieving sensitive detection of low-abundance RNA targets is critical. We evaluated analytical sensitivity by spiking heat-inactivated HIV virions (10–10⁴ copies) into 1 mL plasma-hybridization buffer mixtures containing 5% SDS. Extraction yield was linear across the tested input range and closely followed the standard curve (Fig. 3D). Minor Cq variation was observed at the 10-copy input, likely reflecting stochastic effects associated with low template copy numbers. Occasional apparent recovery exceeding 100% likely reflects quantification variability inherent to RT-qPCR. The lowest detectable input was 10 copies of HIV RNA from 250 µL of plasma, corresponding to 40 copies per mL. This limit can be further lowered by increasing sample input volume, given the high scalability of bead-based extraction.

### Compatibility with whole blood

Extraction from whole blood is more challenging than plasma due to abundant proteins, RNases, and PCR inhibitors, where even minor carryover can reduce RT–qPCR efficiency. Plasma separation is most conveniently achieved by centrifugation, which requires bulky equipment and stable power. Direct RNA extraction from whole blood eliminates plasma separation, reduces target loss, and enables capture of both cell-free and intracellular RNA, potentially improving clinical sensitivity.

To evaluate viral lysis and RNA protection in whole blood, HIV-positive patient plasma was spiked into 125 µL of porcine whole blood. Whole blood samples were pre-diluted twofold with PBS and processed using the same protocol as plasma samples, yielding a final volume of 1 mL. Cq values of the extracted RNA aligned with the standard curve (Fig. 3E). Inputs as low as 10 copies were detected, consistent with effective RNase inactivation by 5% SDS.

To assess lower detection limits in a clinically relevant matrix, we increased the blood input to 1 mL. HIV-positive plasma containing 25 HIV RNA copies was spiked into 1 mL porcine whole blood and incubated for 5 min before SDS addition, then processed using a modified large-volume extraction workflow. Optimization experiments supporting these protocol modifications are shown in cSupplementary Fig. S7. No amplification was observed in NTC, whereas all 8 spiked samples amplified successfully, demonstrating detection at 25 copies/mL in whole blood (Fig. 3F). Given how rapidly blood RNases degrade RNA, recovery after this 5-min exposure indicates that SDS rapidly lysed the viral membrane and protected the released RNA. Additionally, despite the high protein content of blood, we observed no protein gelation or bead aggregation.

To estimate how much background nucleic acid co-elutes with the target, we spiked 1 µg of HIV-negative human genomic DNA into PBS and found that only 0.24% was carried over with the beads (Supplementary Fig. S8). Some background DNA therefore co-elutes with the target, but not at levels that shifted Cq relative to the standard curve. Separately, because capture depends only on sequence complementarity, proviral HIV DNA could also be captured from whole blood, although efficiency is expected to be low without a denaturation step. Together, these results demonstrate that our hybridization capture method is compatible with centrifuge-free workflows and enables highly sensitive RNA detection directly from whole blood.

### Contribution of regional accessibility to capture efficiency

Although secondary structure accessibility is widely considered a critical factor for hybridization efficiency, this consensus mostly stems from studies using short probes (<20 nt) in FISH protocols at low temperatures, such as 46 °C^56,57^, and no accessibility metric has been established for longer probes at elevated temperatures. We therefore evaluated whether it predicts capture efficiency for a 70-nt probe.

We defined accessibility (P_unpaired_) as the mean probability that a contiguous 10-nt window within the capture binding site is fully unpaired, computed *in silico* (RNAplfold) with ±150-nt flanking sequence to preserve local structural context. Scanning the full-length HIV genome showed that accessibility varies widely across regions even at elevated temperature (Supplementary Fig. S4). From this scan, we selected two conserved regions with contrasting predicted accessibility: *env* (gp41; mean P_unpaired_ = 0.012, 1.8th percentile genome-wide) as a poorly accessible target and *pol* (IN; mean P_unpaired_ = 0.220, 76.9^th^ percentile) as a highly accessible target. Published SHAPE reactivity data, an experimental measure of RNA structural accessibility, independently supported the same trend, placing *env* in the bottom 4.2% and *pol* at the 29th percentile genome-wide (Supplementary Fig. S9)^58^. We then measured capture of each region from purified HIV RNA in PBS. Unexpectedly, *env* and *pol* showed comparable recovery both with and without SDS (Fig. 4A) even though *env* is one of the most structured regions in the HIV genome.

**Figure 4.**
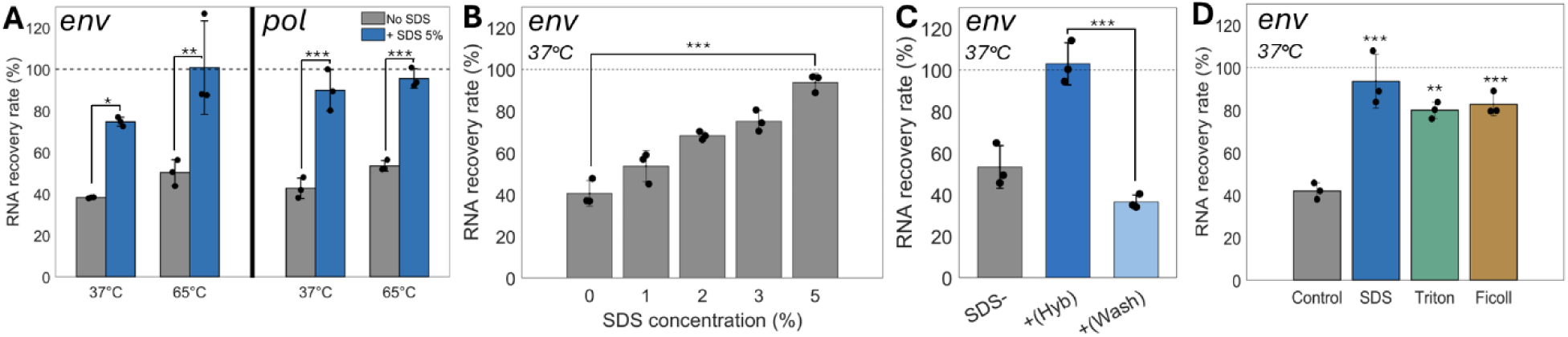
SDS enhances RNA hybridization through a crowding-like mechanism. RNA recovery rate for *env*-and/or *pol*-capture probes, calculated relative to target-matched standard curves (mean ± SD, n = 3, one-way or two-way ANOVA with Tukey’s post-hoc). Purified HIV RNA (10^4^ copies) was spiked into RNase-free PBS. Dashed line indicates 100% recovery. (A) *env* and *pol* at 37 °C and 65 °C, ± 5% SDS. (B) *env* at 37 °C with increasing SDS concentration (0–5%). Recovery increased dose-dependently (Jonckheere–Terpstra trend test), with 0% vs 5% shown. (C) *env* at 37 °C: no SDS (SDS–), SDS during hybridization only (+Hyb), or SDS during a mock wash step only (+Wash), with wash/elution otherwise identical. (D) *env* at 37 °C with no additive (Control), 5% SDS, 5% Triton X-100, or 5% Ficoll 400. Asterisks denote comparisons vs Control.

We speculate that this lack of dependence on accessibility reflects the binding mechanism of long probes. Hybridization initiates through nucleation at short (3-8 nt) unpaired stretches^59,60^, and a 70-nt binding site contains many candidate windows. Once nucleation occurs, the free energy of the 70-nt RNA:DNA duplex overcomes the local target structure, allowing zippering to displace the existing base pairs. In contrast, short probes have fewer candidate windows and dissociate more readily, requiring multiple binding attempts.

Accessibility can also be evaluated as the maximum rather than the mean unpaired probability across the binding site. Under this metric, shortening the window from 10-nt to 3-nt raises both regions but raises *env* more than *pol*, reversing their order (Supplementary Fig. S10). How many such sites are needed, and how long they must be, is not established. Therefore, accessibility was computed for all candidate regions as auxiliary information but was not used for filtering or ranking in the probe design algorithm described below (see Algorithm-guided capture probe design).

### Effect of SDS on hybridization

We initially employed high SDS concentrations to denature proteins and protect RNA from degradation, but our data show that SDS directly enhances RNA hybridization kinetics. Without SDS, raising the temperature from 37 °C to 65 °C increased the recovery to only ∼50% (Fig. 4A). By contrast, pre-heating alone yielded >90% recovery from a synthetic 266-nt RNA target (Fig. 2C), even though the *env*-probe had a more favorable hybridization free energy at 65 °C (ΔG = −54.8 vs. −50.6 kcal/mol, calculated using the NUPACK Python module). This suggests that thermal denaturation alone is not sufficient for every target. Target length may contribute, consistent with Liu *et al*.^61^, who found that pre-heating an intact ∼1,500-nt 16S rRNA target produced minimal gain, whereas fragmenting the target to 20–100 nt increased hybridization efficiency 6-fold.

Adding 5% SDS to PBS dramatically increased recovery across all temperatures. Notably, recovery at 37 °C reached near-saturation, demonstrating that SDS can effectively replace thermal denaturation. SDS titrations revealed a dose-dependent increase in recovery from 0% to 5% (Fig. 4B). In 0.5 M NaCl, the critical micelle concentration (CMC) of SDS is ∼0.05%, suggesting that most of the excess SDS forms micelles that may facilitate hybridization through a molecular crowding effect. To confirm that SDS acts specifically during hybridization rather than during wash, elution, or RT-qPCR steps, we compared adding SDS exclusively during hybridization against adding it only during a mock wash step. Only hybridization-stage SDS enhanced recovery, whereas wash-stage SDS slightly reduced yield relative to the no-SDS control (Fig. 4C). Thus, SDS directly accelerates capture kinetics rather than preventing downstream losses or altering RT-qPCR.

We evaluated two potential mechanisms for this enhancement: surfactant-induced backbone rigidification and macromolecular crowding by abundant micelles. To distinguish these mechanisms, we substituted SDS with either Triton X-100, a non-ionic surfactant above its CMC, or Ficoll 400, a neutral crowding polymer. Both additives restored recovery to levels comparable to SDS (Fig. 4D), supporting macromolecular crowding rather than a unique surfactant chemistry as the primary driver of enhanced hybridization kinetics in protein-free buffer. This crowding mechanism also accounts for the ∼90% recovery observed in plasma with 5% SDS (Fig. 3C). In a protein-rich matrix, SDS partitions into three states: protein–SDS complexes, free monomers, and micelles. Using an established SDS-protein binding stoichiometry (∼1.4g SDS per g protein)^62^ and typical plasma total protein concentration (∼70 g/L), we estimate that approximately 50% of the total SDS would be consumed by protein binding, leaving ∼2.5% free SDS to form micelles. Although 2.5% SDS was insufficient for optimal recovery in PBS (Fig. 4B), plasma containing 5% total SDS still achieved ∼90% recovery. Thus, free SDS alone cannot fully account for the high recovery in plasma. One possibility is that protein–SDS complexes themselves contribute to macromolecular crowding, compensating for the reduced pool of micellar SDS and maintaining efficient hybridization.

### Assessment of RNA Fragmentation after Capture

The capture process involves heating, mixing, and elution steps, all of which could potentially induce RNA fragmentation. If fragmentation occurs during capture and purification, a single-probe system may lose access to long-range genomic information. We therefore spiked synthetic 4-kb RNA into PBS, performed hybridization capture with 5% SDS, and assessed RNA fragmentation by gel electrophoresis (Fig. 5A). The extracted RNA showed a clear band at approximately 4 kb, comparable to that of the non-extracted control RNA in neutralized buffer. This result indicates that short-term heating, bead washing, and pH-based elution did not cause substantial RNA fragmentation. Further, quantification of the 4-kb band intensity was consistent with high recovery by SDS-assisted capture even for this long RNA target (Fig. 5B). Having established that the 4-kb target can be captured intact, we next developed a computational algorithm to select capture probe sequences for clinical detection across diverse HIV subtypes.

**Figure 5.**
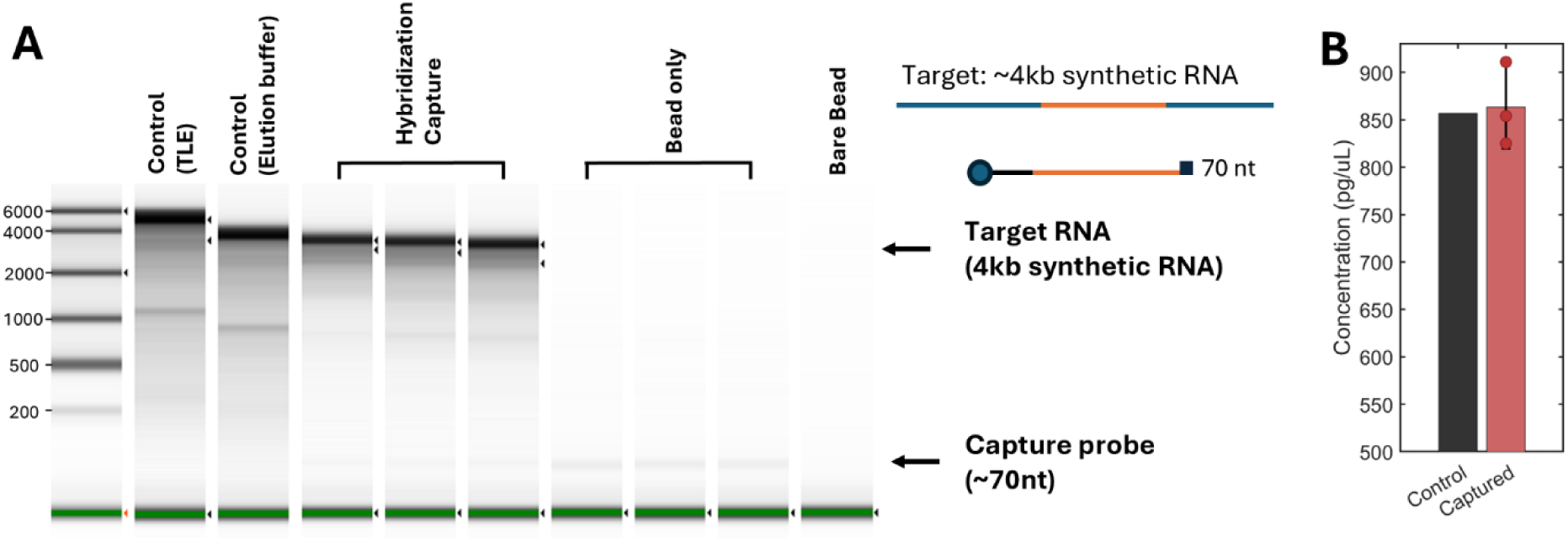
Comparison of RNA fragmentation before and after sequence-specific RNA extraction from PBS with 5% SDS and 70-nt capture probe (*pol* (IN)). (A) ScreenTape analysis of control and extracted RNA.

**Figure 6.**
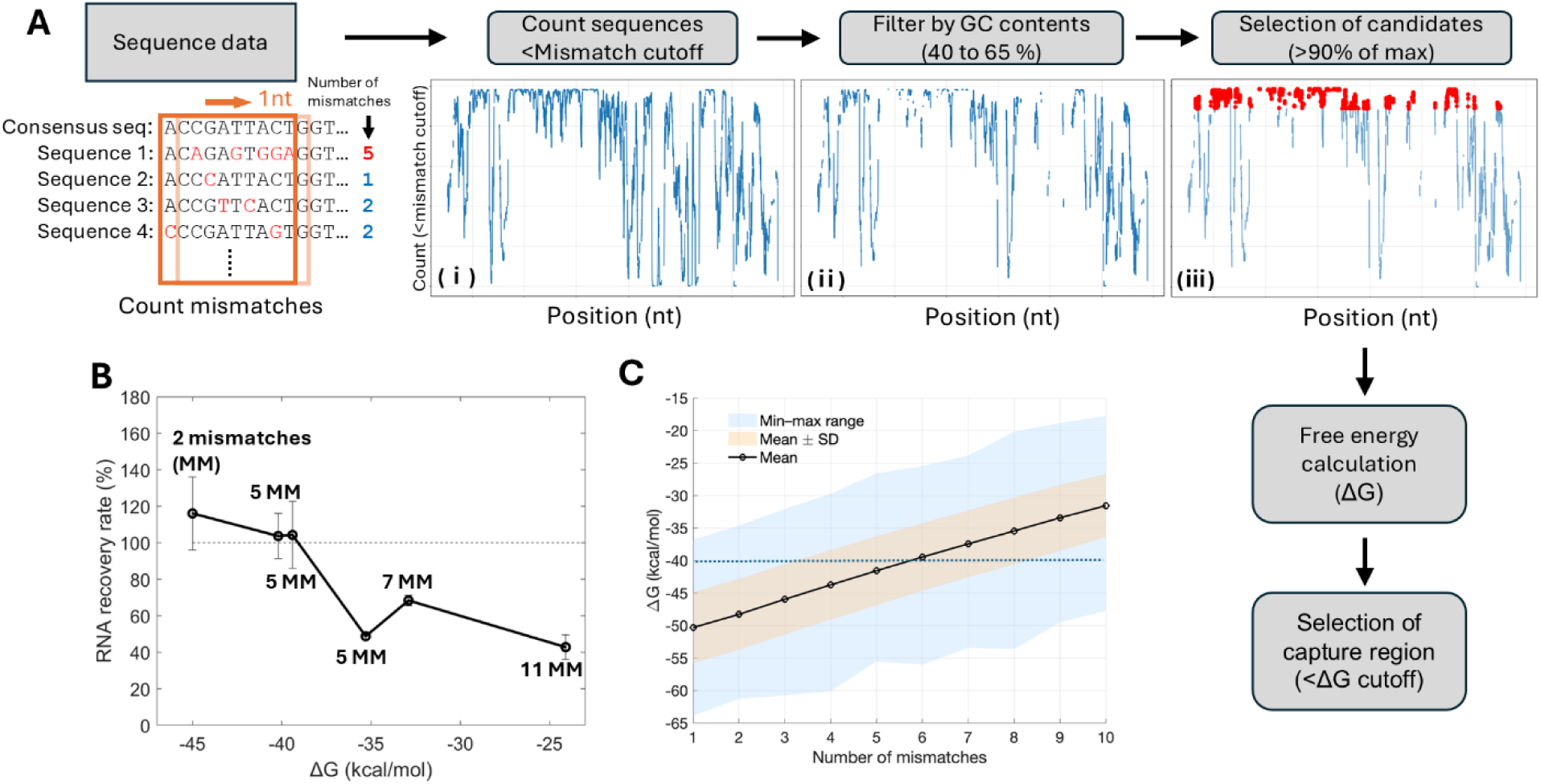
Algorithm for identifying the optimal binding sites. (A) Workflow diagram of the algorithm. (ⅰ) The count of targets that satisfy the mismatch cutoff. X-axis represents the starting nucleotide position for the window of HIV sequence. (ⅱ) Windows with GC content outside the defined range were removed. (ⅲ) Selection of candidate capture windows for thermodynamic analysis. The threshold was set at 90% of the maximum number to cover most of the peaks. (B) Experimentally measured RNA recovery rate for various mismatched HIV RNA targets (10^3^ copies) in PBS + 5% SDS as a function of hybridization ΔG value at 65 °C [Mean ± SD (n = 3)]. 70-nt capture probe was used and target *gag* region. ΔG were calculated using NUPACK (DNA-DNA). The number of mismatches (MM) between the HIV sequence and the capture probe is indicated next to each data point. (C) Simulation results of ΔG values between randomly generated 70-nt strands with varying numbers of mismatches at 65 °C.

Representative TapeStation traces were compiled from multiple runs. The control and captured samples were run together, whereas the ladder was rerun with the control because it failed in the original run, and the bead-only control was run separately. The primary bands correspond to the 4 kb full-length product. Differences in migration between control lanes reflect different ionic conditions rather than fragmentation. Faint bands around 1 kb likely represent T7-derived truncated RNAs retaining the capture probe-binding site. The ∼70-nt band in hybridization and bead-only lanes suggests partial release of the C3-blocked capture probe during elution, which does not interfere with downstream RT-qPCR. (B) Quantification of the band near 4 kb from ScreenTape electropherograms comparing the extracted RNA with the corresponding non-extracted control (Control n=1, captured n=3).

### Algorithm-guided capture probe design

A major concern of hybridization capture is reduced efficiency due to mismatches, which is particularly problematic for highly variable RNA targets. HIV is an appropriate model for evaluating mismatch effects because it is one of the most genetically diverse viral targets^63^. To address the mismatch issue, we developed an algorithm to identify optimal capture regions that maximize efficiency across diverse HIV sequences while minimizing probe count, ideally enabling a single-probe system. The workflow of the algorithm is illustrated in Fig. 6A, and the mechanism is described in the Materials and Methods section. In short, the algorithm first scanned the HIV alignment to identify binding sites satisfying a user-defined mismatch cutoff. This analysis generated a positional plot highlighting peaks where the greatest number of sequences met the mismatch cutoff (Fig. 6A (ⅰ)). Candidates outside the desired GC content range of 40–65% were then filtered out (Fig. 6A (ⅱ)). A subset of the highest-ranked candidates (≥90% of the maximum count number) was selected for thermodynamic evaluation (Fig. 6A (ⅲ)). Hybridization free energies between the consensus sequence and each target sequence were computed for each candidate site. Sequences with a hybridization Gibbs free energy (ΔG) more negative than the user-defined cutoff (<-40 kcal/mol at 65 °C) were predicted to be capturable. Ultimately, the algorithm generated a ranked list of binding sites with predicted capture coverage and thermodynamic metrics.

### Establishing thermodynamic threshold parameters

A primary requirement for the algorithm is to establish a criterion for deciding whether a capture probe can effectively hybridize to mismatched targets. Simple mismatch counts fail to capture the complexity of duplex stability, as the impact depends not only on the number but also on the position and distribution of mismatches. In contrast, ΔG provides a thermodynamic measure that inherently accounts for these variables. To empirically determine a practical ΔG threshold, various HIV mutants were spiked into PBS and 5% SDS mixture, and captured using the same probe (Fig. 6B). When the ΔG between the target and capture probe was below (more negative than) –39.4 kcal/mol, the recovery rates often maintained about 100%, whereas values above –39.4 kcal/mol showed a marked decrease in recovery. Based on these observations, we set –40 kcal/mol as an operational cutoff for efficient capture.

To minimize the computational burden of thermodynamic analysis, we returned to mismatch heuristic to rapidly screen out unlikely candidates. To establish this mismatch heuristic, we simulated random 70-nt sequences (40–65% GC content) with progressively increasing mismatches and fitted a normal distribution to the resulting ΔG values. The probability of a sequence having a ΔG below -40 kcal/mol dropped to ∼4.2% at 10 mismatches (Fig. 6C). Therefore, we used a 9-mismatch cutoff heuristic for 70-nt probes to reduce the candidate pool while retaining thermodynamically strong probes for further analysis.

### Limitations and applicability of algorithms

There are several limitations to this approach. First, the absence of RNA/DNA parameters in NUPACK likely resulted in an overestimation of actual hybridization energies. Nevertheless, since the ΔG cutoff was empirically selected (Fig. 6B), the algorithm relies on relative rather than absolute ΔG, which remains a reliable predictor of capture efficiency within our dataset. Second, the hybridization ΔG values were calculated for solution-phase hybridization and do not explicitly account for ΔG penalties associated with target capture by immobilized probes. Although the ΔG threshold was empirically derived from experimental data, its accuracy may decrease when probe or target lengths differ substantially from those used in our experiments. Such changes could alter entropic penalties, steric hindrance, and hybridization kinetics. In these cases, the ΔG threshold may require recalibration.

### Comparison and selection of candidate capture probe binding sites

We applied the probe design algorithm to HIV sequences (∼9 kb; HIV-1/SIVcpz, Region: GENOME, Subtype: M group with CRFs, 4,927 sequences, Year: 2021) from the Los Alamos database to identify capture-suitable binding sites across genome-wide variation (Table 1).

**Table 1.**
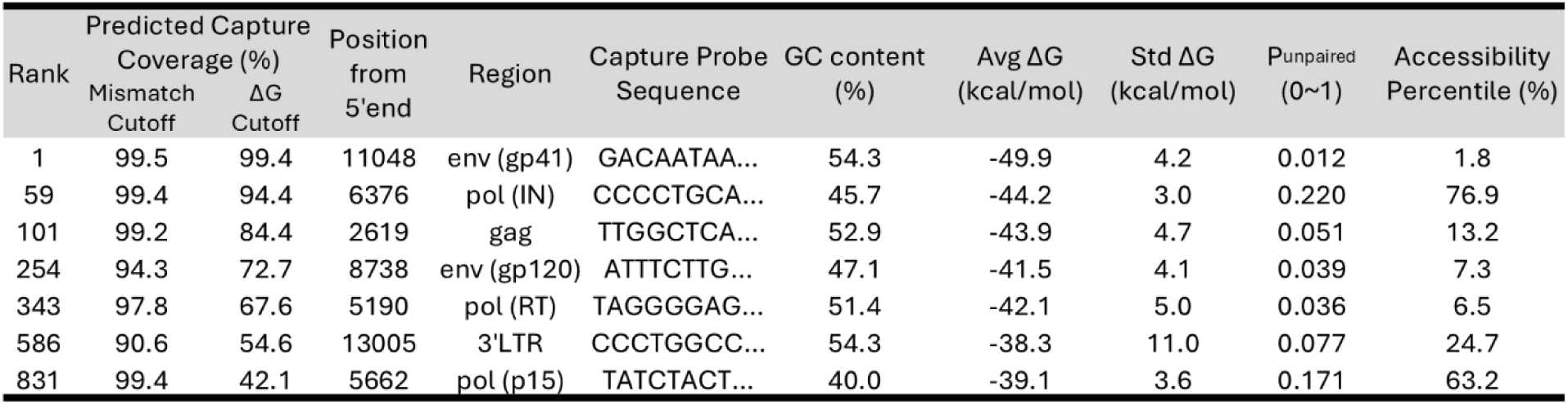
Algorithm-selected 70-nt capture probes based on the HIV-1 M groups with CRFs database. Predicted capture coverage was computed using 2 criteria, a ΔG cutoff and a mismatch cutoff, reported as the percentage of database sequences meeting each cutoff. Regions are ranked by ΔG-based coverage. Only newly identified regions are shown, and missing ranks indicate regions already listed but with small positional shifts. P_unpaired_ is the mean unpaired probability across contiguous 10-nt windows within each binding site at 65 °C. Accessibility percentile represents the relative P_unpaired_ rank of a given 70-nt candidate binding site compared to all possible 70-nt sites across the HIV genome, where higher values indicate greater predicted accessibility.

Across the genome, sites with high predicted capture coverage were primarily concentrated in *env* (gp41), *pol* integrase (IN), and *gag*. Notably, both *env* (gp41) and *pol* (IN) were predicted to achieve >90% capture coverage based on the ΔG cutoff. The relatively permissive mismatch cutoff was intentionally used to avoid excluding potentially suitable targets, resulting in uniformly high mismatch-based coverage estimates. In contrast, applying the more stringent ΔG cutoff substantially reduced predicted coverage for less suitable candidates.

We initially hypothesized that regional accessibility would be an important criterion for probe selection. However, as shown in Fig. 4A, accessibility differences did not meaningfully predict capture efficiency in our system, and we therefore did not use accessibility as a selection criterion. Instead, probe selection was guided primarily by ΔG cutoff-based coverage (Table 1), narrowing the candidates to a small set of high-coverage regions, including *env* (gp41) and *pol* (IN). Although *env* (gp41) showed high predicted capture coverage, its RT-qPCR efficiency was 77%, likely due to its secondary structure. In contrast, the *pol* (IN) region combined high predicted coverage with efficient amplification (>90%). Thus, although regional accessibility did not predict capture efficiency, it may still influence downstream steps such as RT-qPCR amplification. Notably, the *pol* (IN) region is also known to be highly conserved and used as a primer target in clinical RT-qPCR viral load assays^64^, demonstrating robust detection across diverse HIV-1 subtypes. However, to our knowledge, this region has not previously been evaluated as a single capture probe target for sequence-specific RNA extraction. Accordingly, we selected a 70-nt capture probe targeting the *pol* (IN) region.

### Analytical validation using clinical plasma samples

Using a single 70-nt DNA capture probe targeting *pol* (IN), we performed hybridization capture on HIV-positive plasma to evaluate performance in a clinically relevant sample. The HIV-positive plasma sample was diluted with HIV-negative plasma to generate a dilution series.

To assess temperature dependence in clinical samples, Cq values were compared across 25 °C, 37 °C, and 65 °C pre-heating conditions, all in the presence of 5% SDS (Fig. 7A). Cq values differed only modestly across temperatures (25 °C: 32.1 ± 0.20, 37 °C: 32.0 ± 0.17, 65 °C: 31.7 ± 0.12). This difference was not statistically significant (one-way ANOVA, p = 0.058), indicating that heating provides only a modest additional benefit on top of the recovery already achieved by SDS. This is consistent with what we observed for purified RNA in PBS (Fig. 4A). We also noted that below ∼37 °C, SDS–Na^+^ complexes tended to precipitate and aggregate with the beads, complicating handling, whereas the mixture remained fully fluid at ≥37 °C. Because 65 °C avoided this handling issue without any loss in recovery, we retained 65 °C pre-heating as the standard condition for subsequent clinical validation experiments.

**Figure 7.**
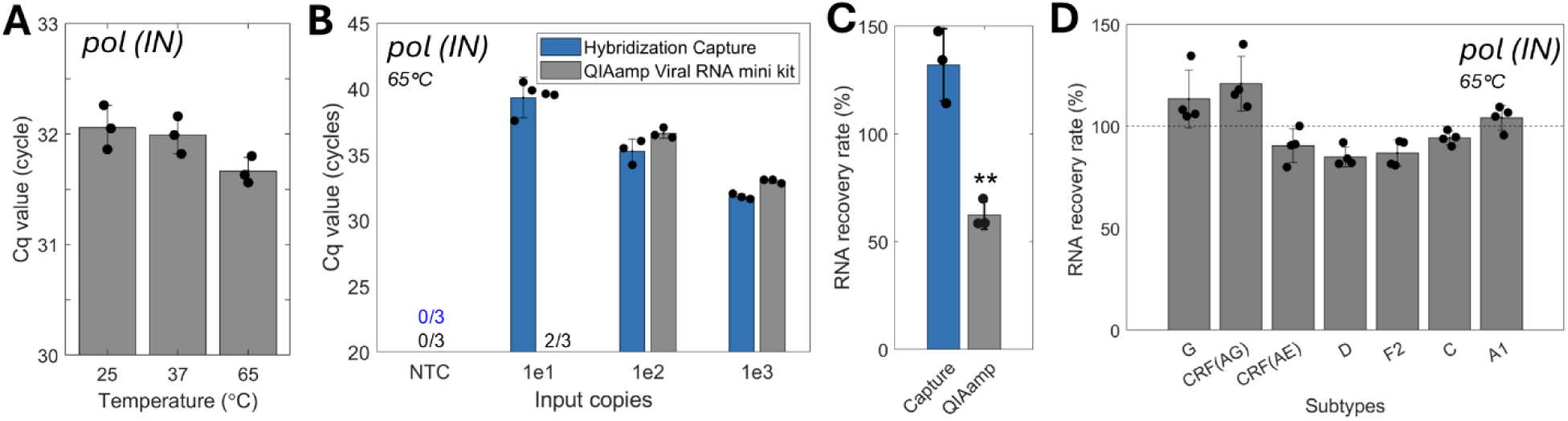
Performance of the algorithm-guided single-probe (70-nt, *pol* (IN)) system across clinical plasma and diverse HIV subtypes with 5% SDS. Panels B–D used 65 °C pre-heating. (A) Cq values for hybridization capture from 140 µL HIV-positive clinical plasma (subtype B, 10³ copies spiked) at 25 °C, 37 °C, and 65 °C (mean ± SD, n=3). (B) Comparison of Cq values obtained with the hybridization capture method and the QIAamp viral RNA mini kit across increasing input copies from 140 µL HIV-positive plasma (subtype B, mean ± SD, n=3). (C) Apparent RNA recovery rate when 10³ copies of HIV RNA were spiked into plasma, comparing capture and QIAamp using purified HIV RNA as a standard curve (mean ± SD, n=3, unpaired t-test). (D) RNA recovery across diverse HIV subtypes using 10⁴ RNA copies from HIV RNA spiked into PBS (mean ± SD, n=4). Recovery rates were calculated based on standard curves generated from 10³ to 10⁵ copies for each subtype.

We next benchmarked the hybridization capture method against the QIAamp Viral RNA Mini kit, a widely used silica-based extraction method, by comparing RNA recovery from clinical plasma via RT-qPCR (Fig. 7B). At the 10-copy input level, the QIAamp kit detected RNA in only two out of three replicates, whereas the hybridization capture method detected all three. At 10²-and 10³-copy inputs, the hybridization capture method also yielded lower Cq values, indicating higher RNA recovery. Because a pooled clinical specimen lacked a matched synthetic standard, apparent recovery rates (Fig. 7C) were calculated against a separate purified RNA stock and should be interpreted with some caution.

To further investigate the QIAamp detection loss noted above, we tested QIAamp recovery in both PBS and clinical plasma across different eluate input volumes (Supplementary Fig. S11). Notably, QIAamp apparent recovery decreased markedly with increasing eluate input volume and was lower in plasma than in PBS. This likely reflects RT-qPCR inhibition by carry-over contaminants. This constraint matters most at low target concentrations, where a large fraction of the eluate must be carried into the reaction. Together, these results highlight the advantage of hybridization capture over silica-based extraction for low-copy RNA detection in complex biological matrices. Finally, to separate the effect of sequence variation from matrix effects, we tested seven HIV subtypes in PBS (Fig. 7D). Mean recovery remained consistently high (>85%) across all subtypes, indicating that the algorithm-designed probe tolerates the mismatches present in these variants.

## Conclusion

Direct RNA hybridization capture from crude samples such as plasma and whole blood has been limited by low recovery and poor reliability. In this study, we quantitatively characterized the effects of temperature, target length, chemical composition, and sample matrix on RNA recovery, and identified conditions that enable highly efficient extraction of target RNA directly from crude samples.

A 266-nt RNA target was recovered efficiently with heating alone, whereas the full-length (∼9 kb) HIV genome was not, likely because heating was insufficient to resolve its extensive secondary structure.

SDS, initially introduced to protect RNA and stabilize plasma by suppressing RNase activity and protein aggregation, had the second benefit of substantially improving recovery of the full-length genome. SDS enhanced hybridization kinetics, potentially through molecular crowding, enabling high RNA recovery without requiring thermal denaturation. SDS-assisted extraction also enabled direct RNA capture from whole blood without centrifugation, detecting HIV RNA at 25 copies per mL. Although this study focused on RNA, the approach may also be applicable to DNA targets (e.g., parasites, bacteria) with appropriate lysis and denaturation conditions.

To design probes that remain robust across HIV subtypes, we developed a ΔG-based probe-design algorithm. Unexpectedly, the mean probability of contiguous 10-nt unpaired windows within the capture site did not predict hybridization performance in our system, even though this metric correctly identified *env* (gp41) as among the most structured regions in the HIV genome. Long probes may overcome local structural barriers through multiple short nucleation sites followed by zippering across the target region. We therefore used an experimentally calibrated ΔG cutoff as the primary criterion for conservative probe selection across HIV sequences. A 70-nt *pol* IN probe designed using this criterion showed robust performance across diverse HIV subtypes and clinical plasma samples.

Direct RNA hybridization capture is particularly suited for enriching low-abundance target RNA from large sample volumes. The method avoids repeated purification and transfer steps that can cause substantial target loss when only a few molecules are present. The bead-based format can be readily scaled with sample volume, allowing rare targets to be concentrated. Combined with compatibility with whole blood without plasma separation, these features also support the development of simplified sample preparation workflows for point-of-care molecular testing, minimizing sample loss during processing. More broadly, the ability to directly recover low-abundance RNA at high efficiency from crude biofluids could overcome a major sample preparation bottleneck in early molecular diagnosis, with potential applications in oncology and the detection of bloodborne pathogens.

## ACKNOWLEDGMENTS

This work was supported by NIH/NIAD as part of a project to develop tests for HIV drug resistance detection (NIH/NIAID R01AI145486. PI: Lutz). We thank all members of the Lutz Lab for their valuable discussions and support. We are especially grateful to Dr. Nuttada Panpradist for initiating this work and for her insightful advice throughout the project. We thank Dr. Ceejay Boyce for providing HIV samples. This work was also supported in part by the acquisition of EQAPOL samples using federal funds from the National Institute of Allergy and Infectious Diseases (NIAID), National Institutes of Health (NIH), under Contract HHSN272201700061C. We thank Greg Pepper and Alex Greninger for facilitating the collection of de-identified remnant plasma samples from the clinical laboratory. We also thank Dr. Xiaohu Gao, Dr. Ayokunle Olanrewaju, and Dr. Wendy Thomas for their helpful advice and discussions.

## Author Contributions

W.K. designed and performed the experiments and wrote the manuscript. S.G. purified and quantified HIV RNA used as controls. E.K. contributed to data analysis and provided constructive input on the experimental design. I.B. characterized and quantified clinical patient samples. L.F. provided feedback and discussion on the manuscript. B.L. contributed to experimental design, manuscript organization, and revision.

## Data availability

The authors declare that all data supporting the findings of this study are available within the paper and its Supplementary information files or available from the corresponding author upon request.

## Code availability

The capture probe design algorithm is available at https://github.com/WookyeomKim/RNA-Capture-Probe-Designer. Custom MATLAB scripts used for thermodynamic profiling, genome-wide accessibility analysis, and ΔG simulation are available from the corresponding author upon request.

