## Supplementary Information for "Surfactant-Enhanced Hybridization Capture for Direct Sequence-Specific RNA Extraction from Plasma and Whole Blood"

### Table of Contents

#### Supplementary Table

**Supplementary Table S1**. Sequences of PCR primers, DNA capture probes, and target RNA.

#### Supplementary Figures

**Supplementary Figure S1.** Custom-designed magnetic rack.

**Supplementary Figure S2.** Predicted MFE structure at 25℃.

**Supplementary Figure S3.** Predicted MFE structure at 65℃.

**Supplementary Figure S4.** Temperature-dependent accessibility and hybridization.

**Supplementary Figure S5.** Compatibility of guanidinium salts.

**Supplementary Figure S6.** Carryover effect on RT-qPCR.

**Supplementary Figure S7.** Optimization of RNA extraction from an 8 mL mixture.

**Supplementary Figure S8**. Minimal background DNA carryover during hybridization capture.

**Supplementary Figure S9.** SHAPE reactivity of the HIV-1 genome.

**Supplementary Figure S10** Accessibility ranking depends on window length.

**Supplementary Figure S11.** Limitation of QIAamp Viral Mini kit.

| Primer | Sequence |
|---|---|
| Synthesis and capture test with 266-nt RNA. [Capture probe length variation test (Figure 2 A-C)] | |
| DNA template (290nt) for synthetic RNA IVT (266nt) | GGGAATTTAATACGACTCACTATAG CAGAGGAGCTCTCTCGACGCAGGACTCGGCTTGCTGAAGCGCGCACGGCAAG AGGCGAGGGGCGGCGACTGGTGAGTACGCCAAAAATTTTGACTAGCGGAGGC TAGAAGGAGAGAGATGGGTGCGAGAGCGTCAGTATTAAGCGGGGAATTTTCT TCAGAGCAGACCAGAGCCAACAGCCCCACCAGAAGAGAGCTTCAGGTCTGGG GTAGAGACAACAACTCCCCCTCAGAAGCAGGAGCCGATAGACAAGGAACTGT ATCCT |
| 19-nt capture probe | /5Biosg/AAAAAAAAAAAAAAAAAAAA CAGTTCCTTGTCTATCGGC/3SpC3/ |
| 40-nt capture probe | /5Biosg/AAAAAAAAAAAAAAAAAAAA TTGTCTATCGGCTCCTGCTTCTGAGGGGGAGTTGTTGTCT/3SpC3/ |
| 70-nt capture probe | /5Biosg/AAAAAAAAAAAAAAAAAAAA AGGATACAGTTCCTTGTCTATCGGCTCCTGCTTCTGAGGGGGAGTTGTTGTCTC TACCCCAGACCTGAAG/3SpC3/ |
| 120-nt capture probe | /5Biosg/AAAAAAAAAAAAAAAAAAAA AGGATACAGTTCCTTGTCTATCGGCTCCTGCTTCTGAGGGGGAGTTGTTGTCTC TACCCCAGACCTGAAGCTCTCTTCTGGTGGGGCTGTTGGCTCTGGTCTGCTCTG AAGAAAATTCCC/3SpC3/ |
| Forward primer | CAGGACTCGGCTTGCTGAA |
| Reverse primer | CGCACCCATCTCTCTCCTTC |
| TaqMan probe | /56-FAM/AGCCTCCGC/ZEN/TAGTCAAAATTTTTGGC/3IABkFQ/ |
| One-step RT-qPCR protocol | [GC-rich reaction buffer]<br>Initial incubation at 55 °C for 6 min and 94 °C for 3 min,<br>followed by 50 cycles of 94 °C for 30 s, 59 °C for 30s, and 68 °C for 1 min. |
| HIV DNA preparation primer | |
| Outer forward primer | AAATCTCTAGCAGTGGCGCCCGAACAG |
| Outer reverse primer | TGAGGGATCTCTAGTTACCAGAGTC |
| Inner forward primer | ACAGGGACCTGAAAGCGAAAG |
| Inner reverse primer | CTAGTTACCAGAGTCACACAACAGACG |
| 4kb truncated HIV RNA synthesis | |
| Forward primer for T7 promoter sequence embedding | GCCCGCTATAAT TAATACGACTCACTATA GGGATAAT TGGAAAGGAAGGACACCAAATGAAAGATTG |
| Secondary forward primer for amplification | GCCCGCTATAATTAATACGACTC |
| Reverse primer | AGTGGTGGTTGCTTCTTTCC |
| PCR protocol | Initial incubation at 94 °C for 30s followed by 15 cycles of 94 °C for 30 s, 61 °C for 15s, and 68 °C for 4min.<br>After gel extraction, 4kb dsDNA was incubated at 50 °C for 1hr with Hi-T7 RNA polymerase |

| 4kb synthetic RNA extraction test (Figure 5) | |
|---|---|
| RNA target | 4kb truncated HIV RNA |
| 70-nt capture probe | /5Biosg/AAAAAAAAAAAAAAAAAAAA CCCCTGCACTGTACCCCCCAATCCCCCCTTTTCTTTTAAAATTGTGAATGAATAC TGCCATTTGTACTGC/3SpC3/ |
| HIV RNA extraction test from plasma (*pol* IN) (Figure 3A-D, Figure 7A-D) | |
| 70-nt capture probe | /5Biosg/AAAAAAAAAAAAAAAAAAAA CCCCTGCACTGTACCCCCCAATCCCCCCTTTTCTTTTAAAATTGTGAATGAATAC TGCCATTTGTACTGC/3SpC3/ |
| Forward primer (80%) | CCHMTTTGGAAAGGACCAGC |
| Forward primer (20%) | CCAGTTTGGAAAGGACCAGC |
| Reverse primer(60%) | CACCTGCCATCTGTTTTCCATA |
| Reverse primer(40%) | CSCCTGCCATCTGTTTTCCATA |
| TaqMan probe | /56-FAM/CTYTGGAAA/ZEN/GGTGAAGGGGCAGT/3IABkFQ/ |
| One-step RT-qPCR protocol | [Standard reaction buffer] Initial incubation at 55 °C for 10 min and 94 °C for 3 min, followed by 50 cycles of 94 °C for 30 s, 54 °C for 30 s, and 68 °C for 1min. |
| HIV RNA extraction test from blood (*pol* IN) (Figure 3E-F) | |
| 70-nt capture probe | /5Biosg/AAAAAAAAAAAAAAAAAAAA CCCCTGCACTGTACCCCCCAATCCCCCCTTTTCTTTTAAAATTGTGAATGAATAC TGCCATTTGTACTGC/3SpC3/ |
| Forward primer (80%) | CCHMTTTGGAAAGGACCAGC |
| Forward primer (20%) | CCAGTTTGGAAAGGACCAGC |
| Reverse primer (100%) | CCTGTCTACYTGCCACACAATC |
| TaqMan probe | /56-FAM/CTYTGGAAA/ZEN/GGTGAAGGGGCAGT/3IABkFQ/ |
| One-step RT-qPCR protocol | [Standard reaction buffer] Initial incubation at 55 °C for 12 min and 94 °C for 30s, followed by 50 cycles of 94 °C for 15 s, 57 °C for 30 s, and 68 °C for 30s. |
| HIV RNA extraction test from PBS (*env* gp41)[Figure 4] | |
| 70-nt capture probe | /5Biosg/AAAAAAAAAAAAAAAAAAAAGACAATAATTGTCTGGCCTGTACCGTC AGCGTTATTGACGCCGCGCCCATAGTGCTTCCTGCTGCTCCCA/3SpC3/ |
| Forward primer | GCTATAGAGGCGCAACAGC |
| Reverse primer | GGCACAGTAGTGGTGCAGATG |
| Intercalating dye (Evagreen) | - |
| One-step RT-qPCR protocol | [GC-rich reaction buffer] Initial incubation at 55 °C for 6 min and 94 °C for 3 mins, followed by 50 cycles of 94 °C for 30 s, 58 °C for 30 s, and 68 °C for 1 min. Melting curve analysis step was included (65 °C to 95 °C, increment 0.5°C) |
| Mismatch analysis to determine ΔG cutoff (*gag*) (Figure 6B) | |
| 70-nt capture probe | /5Biosg/AAAAAAAAAAAAAAAAAAAA AAAGGCCAGGGTTAAGTGCGAATCTTTCCAGCTCCCTGCTTGCCCATACTAAAT GTTTTAGTCTATATTT/3SpC3/ |

| | |
|---|---|
| Forward primer | GAAGGAGAGAGATGGGTGCG |
| Reverse primer | CCCCTGGCCTTAACCGAATT |
| TaqMan probe*<br>(JX140651, KC596062, JX140658) | /56-FAM/AGCGTCAGT/ZEN/ATTAAGCGGGGGAA/3IABkFQ/ |
| Evagreen*<br>(JX140675, KC473845, KC596061) | - |
| Target RNA, leftmost<br>(2 MM, ΔG: -45.0 kcal/mol) | JX140651 |
| Target RNA<br>(5 MM, ΔG: -40.2 kcal/mol) | JX140675 |
| Target RNA<br>(5 MM, ΔG: -39.4 kcal/mol) | KC596062 |
| Target RNA<br>(5 MM, ΔG: -35.3 kcal/mol) | KC473845 |
| Target RNA<br>(7 MM, ΔG: -32.9 kcal/mol) | KC596061 |
| Target RNA, rightmost<br>(11 MM, ΔG: -24.1 kcal/mol) | JX140658 |
| One-step<br>RT-qPCR protocol with Taqman probe<br>(JX140651, KC596062, JX140658) | [GC-rich reaction buffer]<br>Initial incubation at 55 °C for 6 min and 94 °C for 30 s,<br>followed by 45 cycles of 94 °C for 30 s, 53 °C for 30 s, and 68 °C for 1 min. |
| One-step<br>RT-qPCR protocol with Evagreen.<br>(JX140675, KC473845, KC596061) | [GC-rich reaction buffer]<br>Initial incubation at 55 °C for 6 min and 94 °C for 30 s,<br>followed by 45 cycles of 94 °C for 30 s, 55 °C for 30 s, and 68 °C for 1 min.<br>Melting curve analysis step was included (65 °C to 95 °C, increment 0.5°C) |

**Supplementary Table S1.** Sequences of PCR primers, DNA capture probes, and target RNA.

The threshold for Ct determination was set automatically for each run using the Bio-Rad CFX Maestro software based on the baseline fluorescence.

*Given the frequent mismatches in certain regions, RT-qPCR efficiency for some targets was compromised by mismatches in the TaqMan probe region. Consequently, EvaGreen was substituted for these targets alongside minor protocol adjustments. To minimize bias in comparative analysis, all RNA recovery rates were determined based on standard curves processed using identical corresponding protocols.

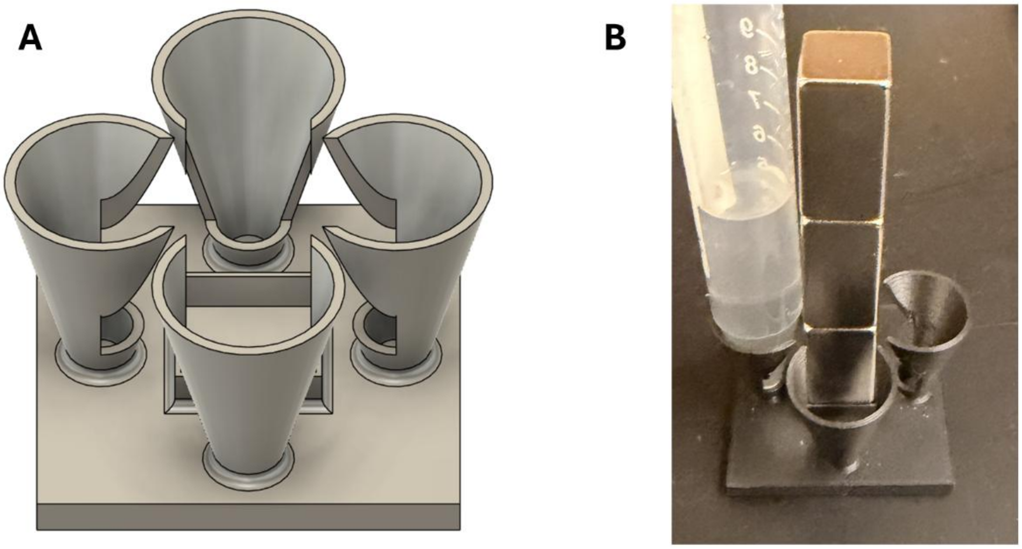


**Supplementary Figure S1.** Custom-designed magnetic rack compatible with 15 mL conical tubes. (A) Three-dimensional design of the magnetic rack. (B) Assembled rack fabricated by 3D printing and integrated with permanent magnetic blocks. The total material cost was approximately USD 15.

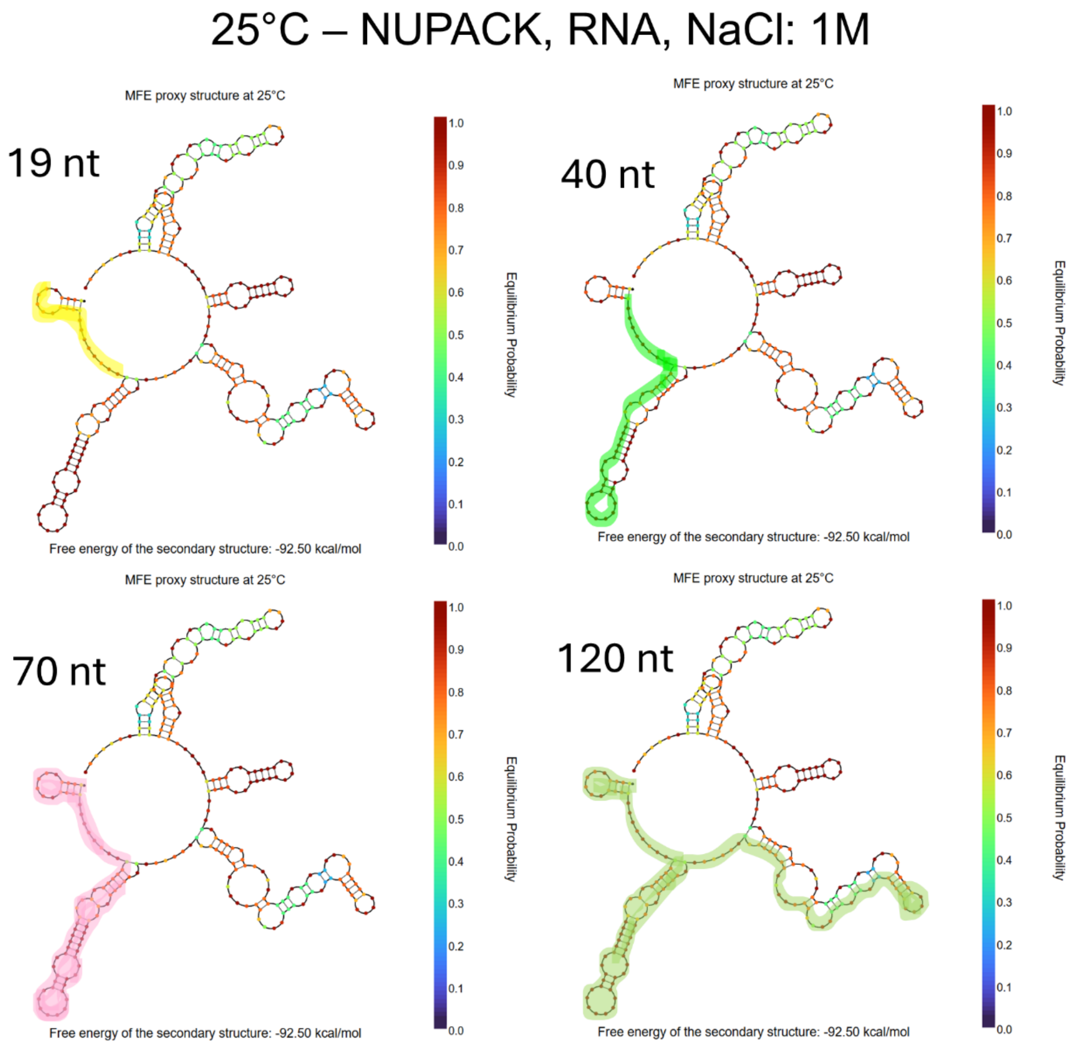


**Supplementary Figure S2.** Predicted MFE structure of the 266 nt synthetic target RNA generated by NUPACK at 25℃. The highlighted regions indicate the binding sites corresponding to each capture probe length. Each binding site was designed with slight positional shifts to maintain a similar GC content of approximately 52 percent.

At 25℃, the target RNA adopts a stable secondary structure that substantially limits the accessibility of the capture probe binding regions. This structure can also displace short capture probes through strand rearrangement, which may together result in a low RNA recovery rate. According to the MFE proxy structure predicted by NUPACK, the 19 nt capture probe binding site appears to be largely free of secondary structure. However, the MFE proxy represents only one of many possible conformations and cannot capture the full structural ensemble of the RNA.

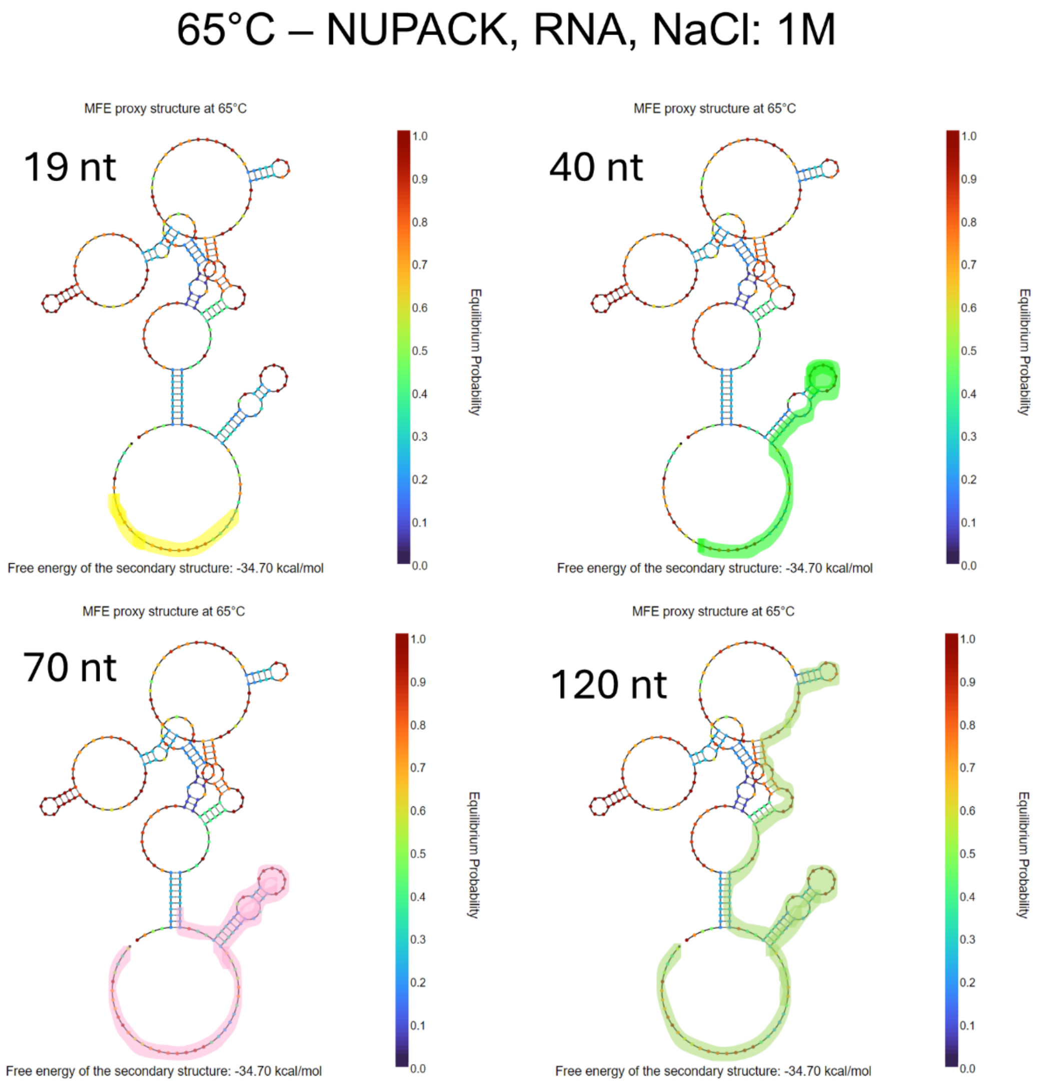


**Supplementary Figure S3**. Predicted MFE structure of the 266 nt synthetic target RNA generated by NUPACK at 65℃. The highlighted regions indicate the binding sites corresponding to each capture probe length.

Most of the stem structures began to unfold at 65℃, and the equilibrium probabilities of many secondary structural elements were noticeably reduced, thereby allowing the capture probes to access and hybridize with the target more efficiently

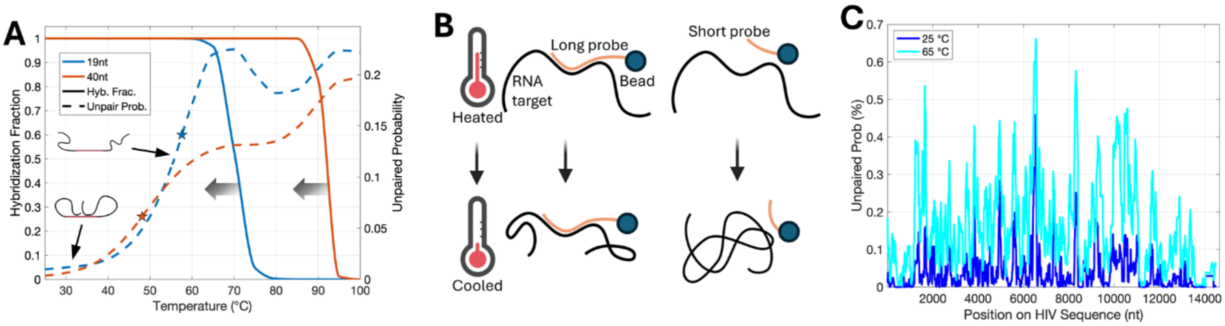


**Supplementary Figure S4.** (A) Temperature-dependent accessibility and hybridization fraction, obtained from *in silico* RNA folding (ViennaRNA) and the RNA-DNA nearest-neighbor model (Sugimoto 1995)[1]. Unpaired probability represents the average unpaired probability across contiguous 10-nt segments within the capture probe binding site. The arrow indicates that bead-based capture may reduce the melting temperature due to a free energy penalty for target hybridization to an immobilized probe (B) Schematic illustration of temperature-mediated capture for short and long probes. (C) Genome-wide $P_{unpaired}$ across the HIV consensus sequence (Los Alamos National Laboratory (LANL), HIV-1/SIVcpz, Region: GENOME, Subtype: ALL, Year: 2021). $P_{unpaired}$ was calculated as the mean probability of contiguous 10-nt unpaired segments within each 70-nt capture window, which was shifted in 10-nt increments across the genome.

We assessed how heating affects target accessibility and probe hybridization by analyzing temperature-dependent accessibility and hybridization fractions (Fig. S4A). Given that hybridization initiates through nucleation on contiguous stretches of unpaired bases[2,3], such segment unpaired probabilities have been widely used to quantify local target availability[4–7]. Accordingly, we defined accessibility as the mean of the probabilities that all possible contiguous 10-nt segments within the binding site are fully unpaired in the structural ensemble.

For the 19-nt and 40-nt capture regions, $P_{unpaired}$ (Fig. S4A, dashes) increased sharply with temperature up to ~65 °C and then reached a modest plateau. Hybridization modeling (Fig. S4A, solid lines) indicates that at 65 °C the 40-nt target-probe complex remains thermodynamically stable, whereas the 19-nt probe is predicted to have a markedly reduced bound fraction. Although the 19-nt probe may rehybridize upon cooling, target accessibility is also expected to decline rapidly with cooling and be kinetically faster than probe hybridization. Under these conditions, probe-target hybridization must compete with rapid intramolecular refolding of the RNA, thereby slowing hybridization kinetics and reducing capture efficiency (Fig. S4B). Together, these results indicate that efficient capture requires not only heat-induced target accessibility, but also probe designs that remain stably bound at the temperatures where the target becomes accessible. Hybridization predictions in Fig. S4A were computed for free solution hybridization, however immobilization on solid surfaces can reduce melting temperatures by up to ~20 °C relative to solution-phase predictions [8,9], which could explain the low capture efficiency of the 19-nt probe (i.e., the blue curve in Fig. S4A overestimates the melt temperature).

Across most regions of the HIV genome, increasing the temperature to 65 °C led to higher $P_{unpaired}$ values (Fig S4C). However, some regions showed little to no increase even at elevated temperature.

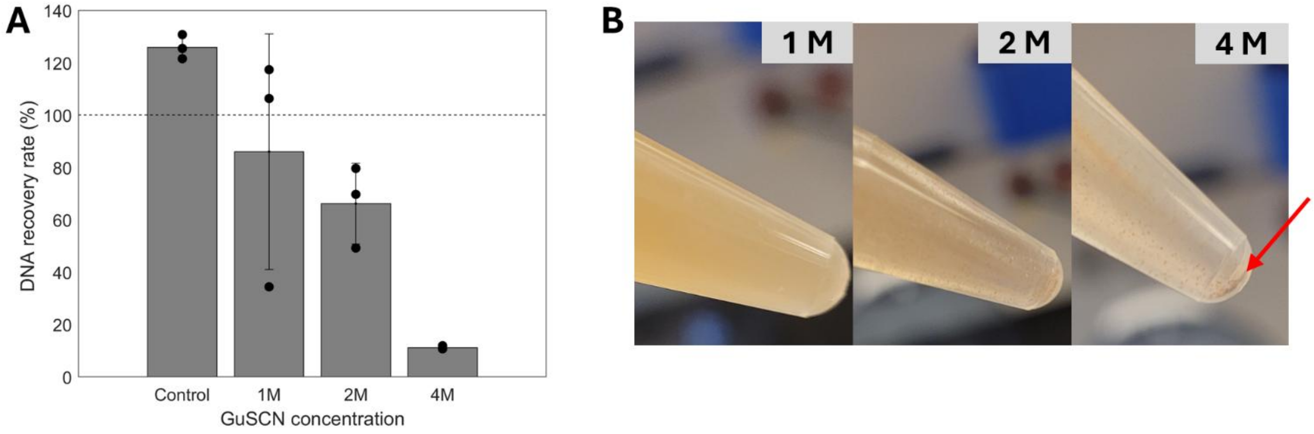


**Supplementary Figure S5.** Compatibility of guanidinium salts with the hybridization capture method without SDS. The quantitative data are presented as mean ± SD (n = 3).
(A) DNA recovery rate as a function of guanidinium thiocyanate (GuSCN) concentration. The template and capture probe used in this experiment were identical to those used in our previous TB cfDNA extraction protocol[10]. (B) High concentrations of GuSCN caused magnetic bead aggregation and precipitation. These results indicate that GuSCN is incompatible with magnetic bead–based extraction.

The experiment followed the hybridization capture protocol described in the main text, except that SDS was replaced with GuSCN at the indicated concentration (1, 2, or 4 M) during hybridization. The control contained neither SDS nor GuSCN. The DNA target and capture probe were identical to those used in our previous TB cfDNA extraction protocol[10], and recovery was quantified by qPCR against a matched standard curve as described therein.

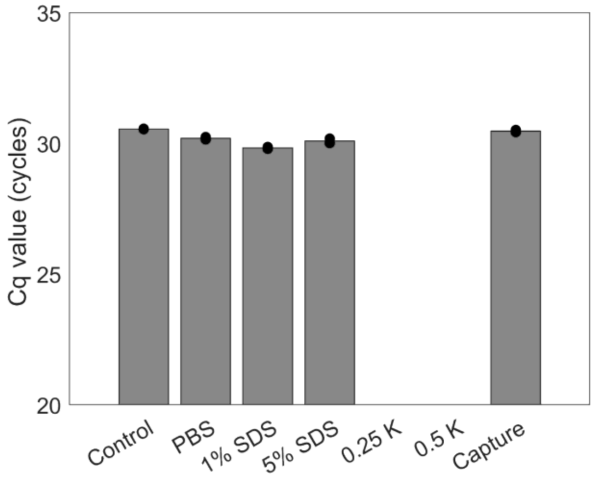


**Supplementary Figure S6.** Carryover effect on RT-qPCR results**.** The quantitative data are presented as mean (n = 2).

To assess whether SDS carryover affects RT-qPCR performance, we conducted a carryover test. The control sample was prepared using freshly made elution buffer without undergoing the hybridization capture procedure. The PBS and SDS groups were prepared using elution buffers collected after the hybridization capture process but without target RNA. For all groups, the elution buffer (without template) was added to the RT-qPCR mixture, followed by the addition of freshly prepared $10^3$ copies of HIV RNA. Negligible differences in Cq values were observed among the control, PBS, and SDS groups, indicating that the washing steps were sufficient to prevent SDS carryover into the RT-qPCR reaction. The *capture* group, which contained RNA obtained through the regular hybridization capture process, also showed similar Cq values, confirming that adding freshly prepared RNA in elution buffer and using extracted RNA yield equivalent amplification efficiency.

Two concentrations of proteinase K were also tested without a heat inactivation step. Normally, proteinase K is inactivated by heating at 95 °C for 10 minutes. However, we were concerned that this high temperature treatment could cause RNA fragmentation or degradation. Therefore, we tested whether residual proteinase K would interfere with the RT-qPCR reaction when the inactivation step was omitted. As shown in the figure, no amplification was observed, indicating that even after three washing steps, the remaining proteinase K was sufficient to inhibit the RT-qPCR reaction.

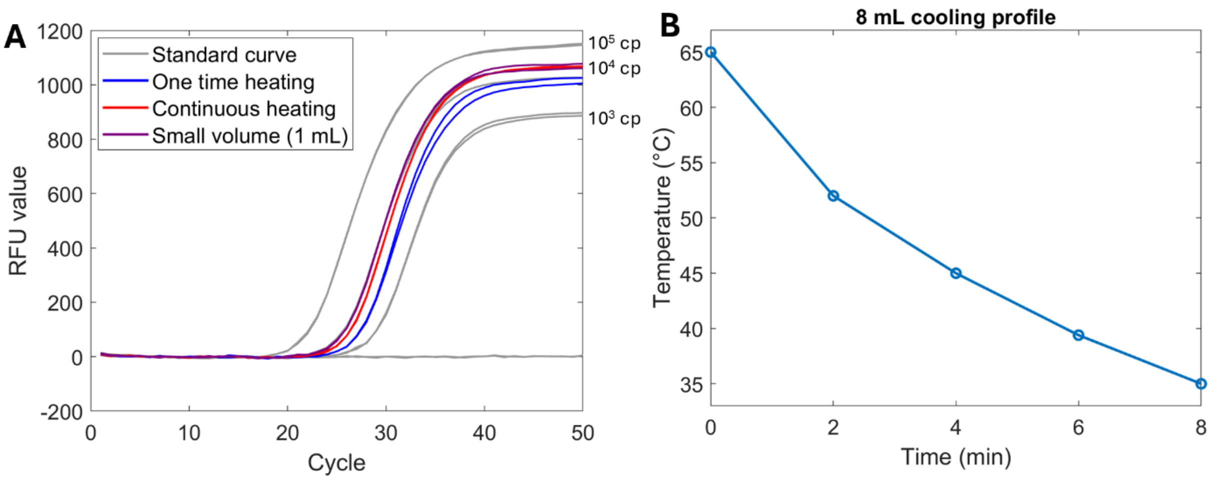


**Supplementary Figure S7.** Optimization of RNA extraction from an 8 mL mixture with 5% SDS. **(A)** RT-qPCR results for HIV RNA extracted from an 8 mL hybridization mixture (PBS). Quantitative data are presented as mean values (n = 2). **(B)** Temperature cooling profile of the 8 mL mixture during rotation (n = 1).

As the sample volume was scaled up from 1 mL of blood to 8 mL, the final hybridization mixture volume increased to 8 mL, necessitating optimization of the pre-heating procedure for efficient RNA extraction. In this experiment, the small-volume (final volume: 1 mL) condition represents the established protocol developed in this work (Fig. S7A). The amplification curves from the small volume mixtures were precisely aligned with the standard curve generated using RT-qPCR master mix and template, indicating efficient HIV RNA extraction from small volume mixtures as we demonstrated throughout this study.

Intuitively, achieving comparable hybridization kinetics for an 8 mL mixture would require proportionally increasing the magnetic bead volume to approximately 160 µL. However, this approach is impractical due to the high cost of magnetic beads. In our previous work, near-quantitative recovery of TB cfDNA from 10 mL urine was achieved using only 50 µL of magnetic beads. Based on this precedent, we optimized the 8 mL extraction protocol using 50 µL of magnetic beads.

In the one-time heating condition, the 8 mL mixture was heated to 65 °C once and then allowed to cool during the hybridization step. Under this condition, the calculated RNA recovery rate was approximately 30%. We hypothesize that hybridization kinetics are substantially slower in the 8 mL volume compared to the 1 mL condition. Although the mixture was initially heated to 65 °C, the temperature dropped to approximately 45 °C within 4 minutes (Fig. S7B). As a result, secondary structures in the target RNA likely reformed before the capture probes could effectively encounter and hybridize with the target sequences, leading to a markedly reduced recovery rate.

In contrast, the continuous heating condition involved incubating the 8 mL mixture in a 70 °C heating block for 15 minutes with periodic tube inversion for mixing, followed by an additional 15 minutes of rotation at room temperature. This continuous heating strategy significantly improved RNA recovery to approximately 65%, nearly doubling the recovery compared to the one-time heating condition. These results suggest that prolonged destabilization of RNA secondary structures is required to allow sufficient time for capture probes to locate and hybridize with target RNA molecules in large-volume samples.

Despite this improvement, the recovery rate did not reach the level observed in the small-volume condition. We attribute this limitation to suboptimal mixing efficiency during heating. Specifically, intermittent manual tube inversion during heating is less effective than continuous mixing using a rotisserie mixer, which likely limits probe–target encounter frequency during the critical hybridization window. Further optimization of simultaneous heating and mixing, as well as extended hybridization time, may further improve RNA recovery in large-volume samples.

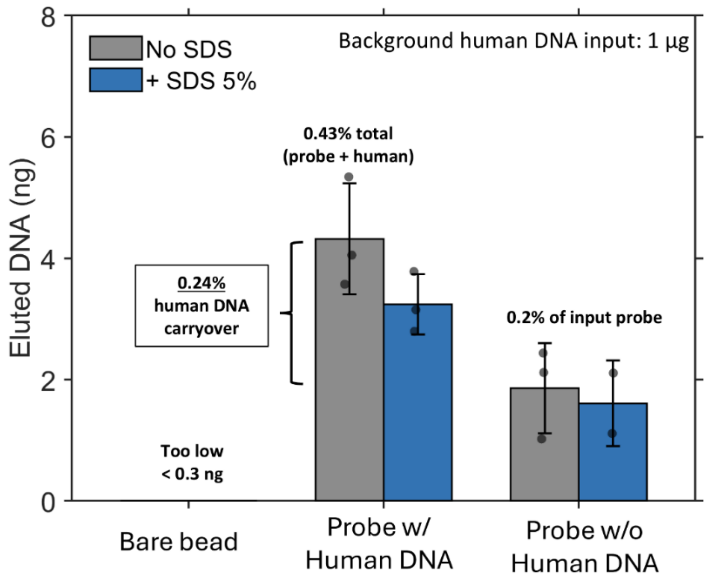


**Supplementary Figure S8. Minimal background DNA carryover during hybridization capture.** Eluted DNA quantified by Qubit ssDNA assay after the standard hybridization procedure in the absence of HIV target, with or without 5% SDS. Bare beads, probe-immobilized beads with 1 µg human genomic DNA spiked into the mixture, and probe-immobilized beads without spiked genomic DNA are compared (mean ± SD, n = 3). Background human DNA input was 1 µg.

To assess the extent of non-specific background DNA carryover, genomic DNA (gDNA) was extracted from human whole blood using the DNeasy Blood & Tissue Kit (QIAGEN, Cat No. 69504). Quantification was performed using the Qubit ssDNA Assay Kit. This kit was selected because the captured target consists predominantly of ssDNA, and the ssDNA assay cross-reacts with dsDNA to reflect total nucleic acid content. A 1 µg aliquot of the prepared gDNA stock (56 ng/µL) was spiked into 1 mL of a PBS–hybridization buffer mixture. Probe-immobilized magnetic beads targeting the HIV pol region were added, and the mixture underwent the standard hybridization process in the absence of HIV targets. Following elution, carryover gDNA was quantified using the same Qubit ssDNA Assay Kit.

While bare magnetic beads exhibited negligible DNA signals, probe-immobilized beads alone (without spiked gDNA) yielded approximately 2 ng of detected DNA in the 23 µL eluate, corresponding to ~0.2% of the input probe amount.

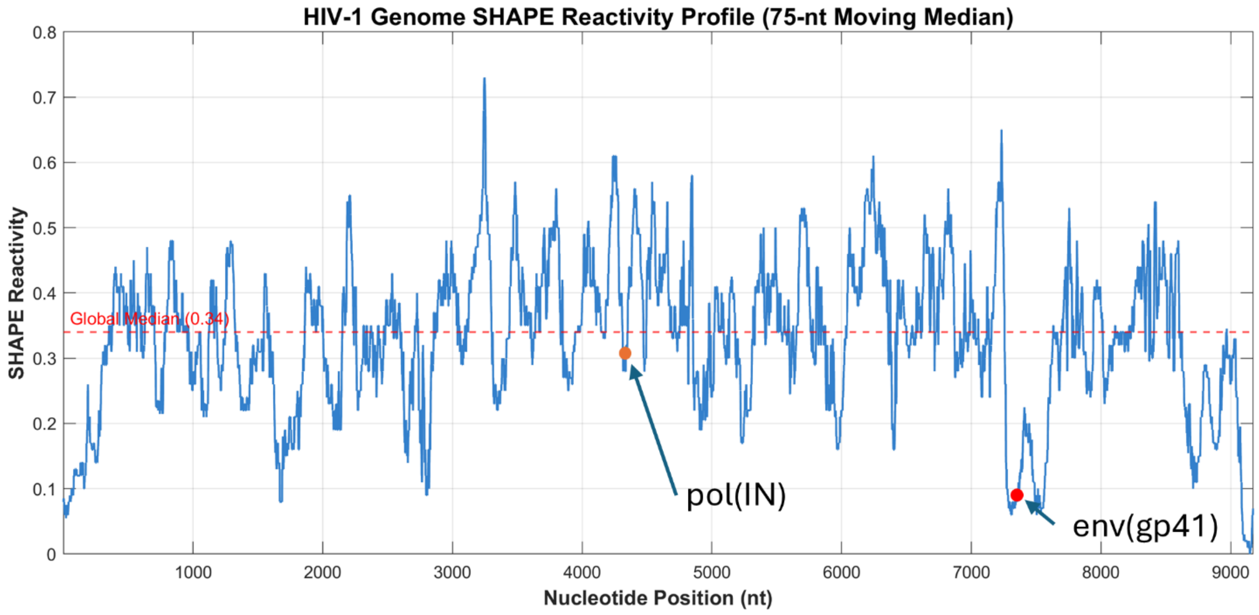


**Supplementary Figure S9. Experimental structure probing places the env capture site among the least reactive regions of the HIV-1 genome.** SHAPE reactivity from [Watts et al., Nature 2009 / dataset][11], shown as a 75-nt moving median to match the scale of the 70-nt capture footprint. The dashed line marks the genome-wide median (0.34). Capture binding sites for pol (IN, orange) and env (gp41, red) are indicated. Percentiles were computed separately from unsmoothed reactivity, taking the median across each 70-nt window genome-wide: env falls at the 4.2nd percentile and pol at the 29th. Capture site coordinates were mapped onto the SHAPE reference sequence by sliding-window sequence matching, selecting the position with minimum Hamming distance.

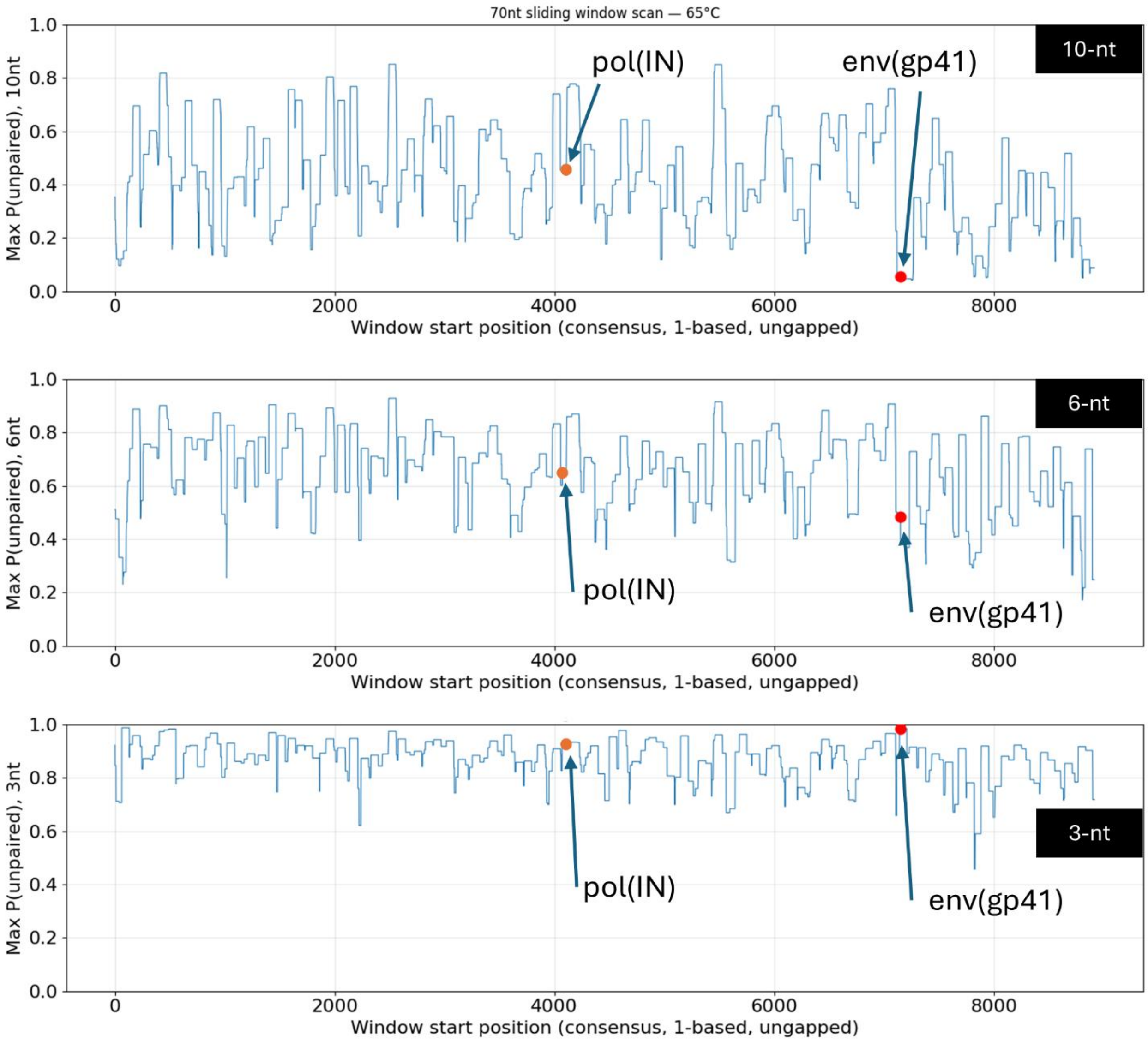


**Supplementary Figure S10. Accessibility ranking depends on the window length used to define an unpaired stretch.** Maximum probability that a contiguous stretch of 10, 6, or 3 nt is unpaired, computed for every 70-nt window across the HIV-1 consensus genome at 65 °C (RNAplfold, ±150 nt flanking). Capture binding sites for pol (IN, orange) and env (gp41, red) are marked. At 10-nt, env is near the bottom of the genome-wide distribution while pol is midrange. The gap narrows at 6-nt and reverses at 3-nt, where nearly all windows in the genome exceed 0.7.

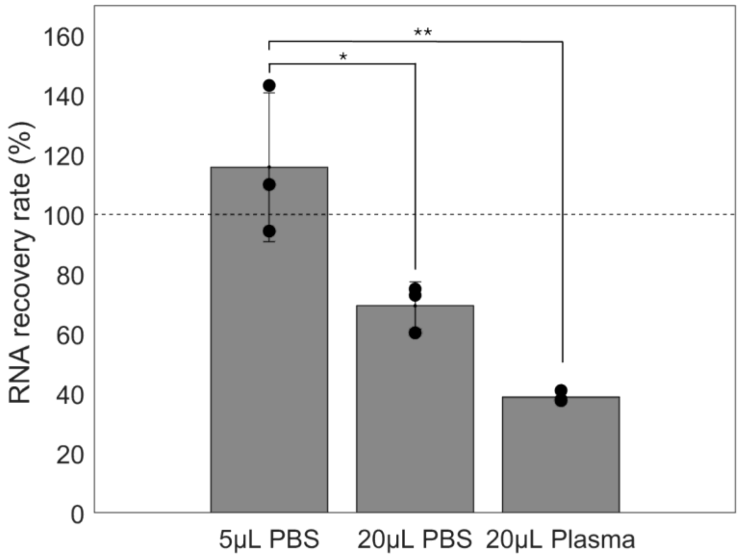


**Supplementary Figure S11**. The limitation of QIAamp Viral Mini kit. The quantitative data are presented as mean ± SD (n = 3).

The QIAamp Viral RNA Mini Kit is considered the gold standard for viral RNA extraction from plasma. Indeed, this kit is a versatile and powerful tool for a wide range of applications, particularly when target sequence information is limited. We evaluated the effect of transfer volume from the QIAamp kit eluate on RT-qPCR performance. Three groups were prepared. In the first group, HIV RNA was extracted from PBS, and 5 µL of the 60 µL eluate was transferred into the RT-qPCR reaction. In the second group, HIV RNA was also extracted from PBS, but 20 µL of the 60 µL eluate was transferred. In the third group, RNA was extracted from a negative plasma sample, and 20 µL of the 60 µL eluate was transferred into the RT-qPCR mixture. A total of $3 \times 10^4$ copies of HIV RNA were spiked into each sample, and recovery rates were calculated based on a standard curve. When only 5 µL of the 60 µL eluate was transferred, the QIAamp RNA recovery rate was close to 100%. In contrast, transferring 20 µL of the eluate significantly reduced the recovery rate to 69.4 ± 8.0%. The eluate may contain residual organic components, such as ethanol from the washing buffer and carrier RNA. While small amounts of these substances may not interfere with amplification, transferring 20 µL into a 50 µL RT-qPCR reaction may be excessive and can adversely affect PCR efficiency. This negative effect was more pronounced when RNA was extracted from plasma samples. Because plasma samples are clinically collected, they contain various genomic materials depending on the patient, which can cause more severe inhibition and result in a much lower recovery rate (38.9 ± 1.86%). These results clearly show that, although the commercial kit is convenient and performs well under ideal conditions, it is not suitable for low-abundance targets.
